# The haplotype-based analysis of *Aegilops tauschii* introgression into hard red winter wheat and its impact on productivity traits

**DOI:** 10.1101/2021.05.29.446303

**Authors:** Moses Nyine, Elina Adhikari, Marshall Clinesmith, Robert Aiken, Bliss Betzen, Wei Wang, Dwight Davidson, Zitong Yu, Yuanwen Guo, Fei He, Alina Akhunova, Katherine W Jordan, Allan K Fritz, Eduard Akhunov

## Abstract

Introgression from wild relatives have a great potential to broaden beneficial allelic diversity available for crop improvement in breeding programs. Here, we assessed the impact of introgression from 21 diverse accessions of *Aegilops tauschii*, the diploid ancestor of the wheat D genome, into six hard red winter wheat cultivars on yield and yield component traits. We used 5.2 million imputed D genome SNPs identified by whole-genome sequencing of parental lines and the sequence-based genotyping of introgression population including 351 BC_1_F_3:5_ lines. Phenotyping data collected from the irrigated and non-irrigated field trials revealed that up to 23% of the introgression lines produce more grain than the parents and check cultivars. Based on sixteen yield stability statistics, the yield of twelve introgression lines (3.4%) was stable across treatments, years and locations; five of these lines were also high yielding, producing 9.8% more grain than the average yield of check cultivars. The most significant SNP-trait and haplotype-trait associations were identified on chromosome arms 2DS and 6DL for spikelet number per spike (SNS), on chromosome arms 2DS, 3DS, 5DS and 7DS for grain length and on chromosome arms 1DL, 2DS, 6DL and 7DS for grain width. Introgression of haplotypes from *Ae. tauschii* parents was associated with increase in SNS, which positively correlated with heading date, whereas haplotypes from hexaploid wheat parents were associated with increased grain width. We show that haplotypes on 2DS associated with increased spikelet number and heading date are linked with multiple introgressed alleles of *Ppd-D1* identified by the whole-genome sequencing of the *Ae. tauschii* parents. While some introgressed haplotypes exhibited significant pleiotropic effects with the direction of effects on the yield component traits being largely consistent with the previously reported trade-offs, there were haplotype combinations associated with the positive trends in yield. The characterized repertoire of the introgressed haplotypes derived from *Ae. tauschii* accessions with the combined positive effects on yield and yield components traits in elite germplasm provides a valuable source of alleles for improving the productivity of winter wheat by optimizing the contribution of component traits to yield.

## 1 Introduction

The gap between population expansion and food production is increasing due to marginal improvements in crop yield, which is attributed to declining soil fertility, pests and diseases, and climate change (Bailey-Serres et al., 2019). Wild relatives of wheat are a rich source of novel underutilized allelic diversity with great potential to improve cultivated wheat through introgression (Placido et al., 2013; Zhang et al., 2017; Hao et al., 2020). Introgression from wild relatives into elite wheat cultivars was reported to increase pest and disease resistance (Periyannan et al., 2013; Saintenic et al., 2013), improve resilience towards environmental stress (Peleg et al., 2005; Placido et al., 2013) and increase yield (Pasquariello et al., 2019). The success of introgression breeding, however, could be affected by the negative epistasis between the multiple alleles of wild and cultivated wheat (Nyine et al., 2020), especially in the low recombing regions of chromosomes, where linkage with the negatively selected alleles could reduce the efficiency of selection for beneficial variants (Hill and Robertson 1966).

Introgression could exhibit pleotropic effects, affecting multiple, often unrelated traits not directly targeted by selection. For example, introgression from *Ae. ventricosa* into chromosome 7D of wheat was associated with increase in grain protein content and resistance to eyespot at the expense of reduced yield (Pasquariello et al., 2019). In durum wheat, introgression of the *GNI-A1* gene from wild emmer increased grain weight by suppressing the fertility of distal florets, resulting in a negative correlation between grain number and grain weight (Golan et al., 2019). Introgression from *Agropyron elongatum* into the 7DL chromosome arm of wheat that is known to confer leaf rust resistance (*Lr19*) (Wang and Zhang 1996) also influences root development, resulting in improved adaption to water stress (Placido et al., 2013) and salinity (Dvorak et al., 1988), and increased biomass and yield (Reynold et al., 2001).

Crop yield is a complex trait determined by many component traits, such as thousand gain weight, grain number per spike, spikes per unit area, grain width, area and length among others (Del Moral et al., 2003; Du et al., 2020; Zhang et al., 2018). Previous studies have shown that introgression from wild and close relatives improve yield of hexaploid wheat by changing yield component traits through the pleotropic interaction between introgressed and background alleles of the hexaploid wheat (Jones et al., 2020). Significant trade-offs between yield, yield components and yield stability have been reported in wheat. A study by Pennacchi et al. (2019) showed that yield and yield stability have a negative linear relationship in most cases. Other factors such as heading date, plant height and biomass influence the source-sink ratio, which in turn affects the harvest index leading to variation in yield and yield stability (Reynold et al., 2001; Reynold et al., 2020). Balancing the trade-off between yield components is therefore necessary to improve yield, maximize the yield potential and improve yield stability in wheat.

Sequence-based genotyping generates high-density SNP marker data that could be used to accurately detect the boundaries of genomic segments introgressed from wild relatives (Kuzay et al., 2019; Nyine et al., 2020), providing a unique opportunity to investigate the effect of introgression on trade-off between traits affecting total yield. Even though whole genome sequencing became less expensive, it is still not within the cost range that would allow wheat breeding programs to sequence large populations. Sequencing of founder lines at a high coverage depth and using these genotypes as a reference panel to impute missing and ungenotyped markers in the progeny characterized by low-coverage SKIM sequencing is a cost-effective alternative. The existing imputation algorithms (Browning and Browning 2013; Swarts et al., 2014; Davies et al., 2016) provide highly accurate whole-genome prediction of missing genotypes, and were shown to increase the power of genome-wide association scans, thus enabling the identification of SNPs or haplotypes associated with the traits of interest (Li et al., 2010; Nyine et al., 2019). One of the advantages of the increased marker density provided by whole-genome sequencing is the ability to perform association mapping using haplotype information, which improves the detection of quantitative trait loci that would otherwise be missed when using single SNPs (Zhang et al., 2002; Lorenz et al., 2010).

Here we investigated the impact of introgression from *Ae. tauschii* into hard red winter wheat lines on yield and yield component traits and how haplotypes from *Ae. tauschii* at different genomic loci affect the component traits and total yield. For this purpose, we assessed the phenotypic variability of yield and the component traits, biomass and tenacious glume traits in an introgression population derived from *Ae. tauschii* and hexaploid winter wheat phenotyped under irrigated and rainfed conditions. We applied SNP diversity data generated by the whole genome shotgun sequencing at 10x coverage level of the parental lines to impute genotypes in this population (Nyine et al., 2020). This strategy resulted in 5.2 million SNPs that enabled us to identify the introgressed *Ae. tauschii* haplotypes and assess their effects on trait variation through genome-wide association mapping and haplotype block effect analysis. This introgression population along with high-density SNP genotype data provides a valuable resource for effective deployment of *Ae. tauschii* haplotypes in winter wheat improvement programs.

## 2 Materials and Methods

### 2.1 Plant materials

The study population was described in detail by Nyine et al. (2020). In brief, the population was developed by crossing synthetic *Ae. tauschii*-wheat octoploid lines with recurrent hexaploid winter wheat lines. The octoploid lines were developed by crossing five hexaploid winter wheat lines with 21 diverse *Ae. tauschii* accessions. The resulting F_1_ hybrid plants regenerated from rescued embryos were treated with colchicine to produce the synthetic octoploids (Dale et al., 2017). The synthetic octoploids were backcrossed once to the respective hexaploid wheat parents or to another wheat line. The BC_1_F_1_ plants (hexaploid) were self-pollinated and advanced by single seed descent to the BC_1_F_3_ generation. Seeds from individual BC_1_F_3_ plants were bulked and grown in single rows in the field at the Kansas State University, Ashland Research Farm near Manhattan, KS in the 2016-17 growing season. A total of 351 lines were selected from the entire population based on the ability to produce sufficient seeds to allow for yield testing, general fitness, threshability and phenology corresponding to the adapted wheat cultivars.

### 2.2 Field phenotyping

The population was phenotyped in 2018 and 2019 at Colby (KS), and in 2020 at Ashland (KS). In both locations and years, an augmented design was used to establish the trials. Plots were planted using a New Holland 6-row wheat drill. Plot dimensions were 2.5 meters long by 0.5 meters wide with 18 cm row spacing. Starter fertilizer was applied with the seed at planting using granular 18-46-0 diammonium phosphate (DAP) at a rate of 168.1 kg/ha. Additional nitrogen was applied as a topdress in the spring using liquid 28-0-0 urea ammonium nitrate (UAN) at a rate of 67.3 kg/ha. A lateral irrigation system was used at Colby to ensure uniform germination and emergence as well as to provide additional water throughout the growing season in the irrigated treatment. Three hexaploid winter wheat lines well-adapted to Kansas environments (checks) and the hexaploid wheat parents were used as controls with at least three biological replications per block. In 2018 and 2019, two complete blocks were established and one block was irrigated (COI18 and COI19, respectively) whereas the other was rainfed/non-irrigated (CO18 and CO19, respectively), simulating an optimal and farmer-field growth conditions. In 2020, only one block was grown under rainfed conditions (AS20).

The population was phenotyped for yield and yield components traits, biomass traits and tenacious glume. Agrobase software (Mulitze 1990) was used to adjust the grain yield (GY), (bushels per acre) for spatial variability. The MARVIN seed imaging system (GTA Sensorik GmbH, Germany) was used to assess the grain morphometric traits such as grain number (GN) per sample, thousand grain weight (TGW), grain area (GA), grain width (GW) and grain length (GL) from two technical replicates in 2018, and one measurement in 2019 and 2020. In 2019 and 2020, data were collected for the number of spikes per square foot (SPSF) from two random points within a plot. The 1 ft x 1 ft square frame was dropped over two rows at least one foot away from the plot edges to avoid the border effect. In 2019, only one row within a frame was cut above the ground level for biomass determination, while in 2020, both rows were sampled. Biomass samples were collected in paper bags and dried for at least three weeks at 32°C (90°F) before processing. We collected data on aboveground dry biomass (BM) measured as the total weight of the dry sample without the bag, the number of spikes per sample (SPB), the average spikelet number per spike (SNS) from 10 random spikes and grain weight after threshing (GSW). During threshing, we scored samples for presence and absence of the tenacious glume (Tg) trait depending on the threshability. Harvesting index (HI) was calculated as the percentage of GSW relative to BM.

In 2020, data for heading date (HD) were collected from each plot when approximately 50% of the spikes had emerged from the flag leaves. The number of days to heading were calculated as the difference between the heading and planting dates. After all the plots had completed heading, plant height (PH), in centimeters was measured on the same day from two random but representative main tillers per plot for the whole field. PH was measured as the distance from the ground surface to the first spikelet of the spike.

### 2.3 Genotyping

#### 2.3.1 Whole genome shotgun sequencing of parental lines

Genomic DNA of 21 *Ae. tauschii* accessions and six hexaploid parents used to create the introgression population was extracted from leaf tissues of two-week old seedlings grown in a greenhouse using DNeasy Plant Mini Kit (Qiagen) following the manufacturer’s protocol. The quality and concentration of the DNA was assessed using PicoGreen dsDNA assay kit (Life Technologies).

Genomic libraries for Illumina sequencing were constructed from ∼2 μg of genomic DNA using the PCR-free Illumina protocol at the K-State Integrated Genomics Facility (IGF). The libraries were subjected to size selection using the Pippin Prep system (Sage Scientific) to enrich for 400-600 bp fragments. The pooled barcoded libraries were sequenced using the NovaSeq instrument (2 × 150 bp run, S4 flow cell) at Kansas University Medical Center and NextSeq 500 (2 × 150 bp run) at IGF. The PCR-free whole genome shotgun sequencing libraries generated from 27 parental lines ranged between 200 and 700 bp with an average of 450 bp (Figure S1). Approximately 14 billion paired-end reads (150 bp long) were generated from the libraries with an average of 0.54 billion reads per sample (data is available at NCBI SRA database BioPrject ID: PRJNA745927). The average number of reads per wheat line corresponded to approximately 10x genome coverage of parental lines. The raw reads with Phred quality score less than 15, minimum length less than 50 bp and adaptor sequencies were filtered out using Trimmomatic v0.38-Java-1.8. The remaining filtered paired-end reads were mapped to the Chinese Spring (CS) RefSeq v1.0 (IWGSC, 2018) using BWA-mem software v0.7.17. A total of 7.1 billion reads were aligned uniquely to the CS RefSeq v1.0.

The sam files generated by BWA-mem were converted to bam files using SAMtools v1.10. Picard Toolkit (http://broadinstitute.github.io/picard) was used to merge bam files from different lanes and sequencers into one bam file per sample. Reads that aligned to multiple locations within the genome were identified and removed by SAMtools v1.10. Picard Toolkit was used to prepare the merged unique aligned read bam files for GATK (McKenna et al., 2010) analysis. The preparatory steps included sorting, adding read groups, marking and removing duplicate reads. The output deduplicated bam files were realigned around INDELs using GATK v3.7 and recalibrated with 90K SNPs (Wang et al., 2014) mapped to CS RefSeq v1.0 as the reference coordinates. The bam files were split into chromosome parts and indexed to reduce the memory and time required to process the files. GATK v3.7 was used to generate the genome variant call format (gvcf) files for each chromosome part. The gvcf files were split into 100 Mb chromosome windows and stored as genomicsDB using genomicsDBImport tool in GATK v4.0. Joint genotyping of variants from each database was done using GATK v4.0 HaplotypeCaller. The flag <-allow-old-rms-mapping-quality-annotation-data> was set to enable the processing of gvcf files generated by GATK v3.7. All vcf files corresponding to the A, B and D genomes were concatenated with concat, a Perl-based utility in vcftools v0.1.13. A custom Perl script was used to convert the Chinese Spring RefSeq v1.0 parts coordinates in the concatenated vcf to full coordinates after which, vcf-filter tools v0.1.13 were used to remove INDELs, multi allelic loci, sites with missing data and MAF below 0.05. The filtered SNPs were phased using Beagle software v4.1 (Browning and Browning 2013).

The GATK v4.0 HaplotypeCaller identified ∼99 million variants including SNPs and INDELs from reads uniquely aligned to the D genome of CS RefSeq v1.0. After excluding INDELs, multi-allelic loci, sites with missing data and MAF below 0.05, 20 million SNPs were retained. These were used to impute missing and ungenotyped SNPs in the D genome of the introgression population.

#### 2.3.2 Genotype imputation

Sequence-based genotyping of the introgression population was performed previously by Nyine et al. (2020). SNPs with minor allele frequency (MAF) less than 0.01 were excluded from the raw vcf file using vcf-filter tools v0.1.13. The program conform-gt (https://faculty.washington.edu/browning/conform-gt.html) was used to check the concordance of D genome SNP positions between the introgression population and the SNPs from parental lines genotyped by whole genome shotgun sequencing. Missing and ungenotyped SNPs in the D genome of the introgression population were imputed from the parental lines using Beagle v.5.0. A custom Perl script was used to filter out imputed SNPs with genotype probability below 0.7 and MAF less than 0.05 which resulted in 5.2 million SNPs. The filtered SNPs were used in the downstream analyses such as genome-wide association mapping and identification of the introgressed haplotype blocks.

### 2.4 Introgressed haplotype detection

Genetic divergence between the parental lines affects the probability of accurate detection of introgressed segments from wild relatives. We used two introgression families, one created by crossing hexaploid wheat with *Ae. tauschii* ssp. *strangulata* (KanMark x TA1642, aka FAM93) and another one created by crossing hexaploid wheat with *Ae. tauschii* ssp. *tauschii* (Danby x TA2388, aka FAM97) to identify introgressed haplotype blocks. FAM93 had 21 introgression lines while FAM97 had 23 introgression lines. The R package HaploBlocker (Pook et al., 2019) was used to infer haplotype blocks from 5.2 million SNP sites. Recombinant inbred lines from each family were analyzed together with 21 *Ae. tauschii* and six hexaploid parents. The HaploBlocker parameters used in block calculation were node_min = 2 (default is 5), overlap_remove = TRUE, bp_map, equal_remove=TRUE. The parameter node_min was used to control the number of haplotypes per node during cluster-merging step of the block calculation function of HaploBlocker. The reduction in node_min was necessary to account for the low number of haplotype variants within these families. To maintain the SNP position in the haplotype block library, a vector of SNPs was provided via the parameter bp_map and prior to block calculation, SNPs in perfect linkage disequilibrium were removed by setting the parameter equal_remove = TRUE. Overlapping haplotypes were removed by setting parameter overlap_remove = TRUE. Custom R and Perl scripts were used to calculate the haplotype block length using the information from haplotype block start and end coordinates. All monomorphic haplotypes between the two parental lines were excluded from the haplotype matrix before calculating the frequency of introgressed haplotypes per chromosome.

### 2.5 Phenotype data analysis

#### 2.5.1 Trait stability and heritability

Yield stability is an important trait, which reflects the productivity of the crop under various growth conditions. No single statistic, however, is accurate enough to predict it due to high variability of genotype and genotype by environment interaction effects. In this study, we used 16 different statistics including parametric (such as Mean variance component (*θi*), GE variance component (*θ*_*(i)*_), Wricke’s ecovalence 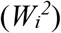, Regression coefficient (*\*b*_*i*_), Deviation from regression (*S*^*2*^_*di*_), Shukla’s stability variance (*s*^*2*^_*i*_), Coefficient of variance (*CVi*) and Kang’s rank-sum (*Kang* or *KR*)) and non-parametric (such as Huhn’s and Nassar and Huhn’s (*S*^*(1)*^, *S*^*(2)*^, *S*^*(3)*^, *S*^*(4)*^, *S*^*(5)*^ and *S*^*(6)*^) and Thennarasu’s (*NP*^*(1)*^, *NP*^*(2)*^, *NP*^*(3)*^ and *NP*^*(4)*^)) methods to rank the introgression lines for yield stability based on their performance across years, locations and treatments. The description and properties of the statistics are documented at: https://manzik.com/stabilitysoft/. The analysis was implemented in R using a script from Pour-Aboughadareh et al. (2019) which is available at: https://github.com/pour-aboughadareh/stabilitysoft. The most stable and/or high yielding lines were identified by sorting them according to their rankings.

Broad sense heritability (*H*^2^) and best linear unbiased predictions (BLUPs) for yield and the component traits were calculated from 2018 and 2019 data. A mixed linear model with restricted maximum likelihood implemented in R package *lme4* was used to generate the variance components (var) that were used to calculate heritability as follows.

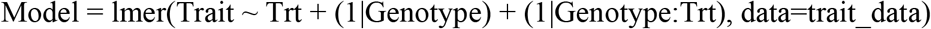

H^2^ = var(Genotype)/[var(Genotype) + var(Genotype:Trt)/No. of Trt + var(Residual)/No. of Trt)]. Where Trt refers to irrigated and non-irrigated treatments. BLUPs were extracted from the linear mixed model as the random effects of genotypes.

Multiple comparison for the effect of treatment on yield and yield components traits in the introgression population relative to the controls was performed using least squares (LS) means with ‘tukey’ adjustment method and α = 0.05. To further assess the impact of introgressed haplotypes on the traits, lines were sorted in a descending order for each treatment and location/year. The percentage of the introgression lines that performed better than the best parental lines and checks (PTPL) was calculated for each trait. Similarly, introgression lines that produced more grains than both parents and checks were identified based on mean spatial adjusted yield. The percentage increase in yield was calculated as follows.

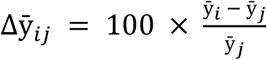

Where 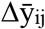 is the percentage change in mean yield, 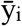 is the mean yield for the high yielding introgression lines and 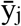 is the mean yield for the controls (parents and checks).

### 2.6 Trade-off between traits

The relationship between yield, yield components and biomass traits were assessed by calculating the Pearson’s correlation coefficients. We compared the trend of correlations from different treatments (irrigated/non-irrigated) and years to determine the extent of trade-off between traits within the introgression population and how the environment influenced them.

### 2.7 Genome-wide association mapping

Genome-wide association (GWAS) analysis was used to identify genomic regions with SNPs and haplotypes that have significant association with the traits. We tested the association of 5.2 million SNPs from the D genome with traits phenotyped in different treatments and years. A total of 15,967 haplotype block windows was identified from 5.2 million SNPs using the R package HaploBlocker v1.5.2 (Pook et al., 2019). Default parameters for HaploBlocker were used except node_min, which was reduced to 2 (default is 5) since most genomic intervals in our dataset had less than 5 haplotype variants. Overlapping haplotypes were removed using the parameter overlap_remove = TRUE and the SNP coordinates were included in the haplotype library via the parameter bp_map. Haplotype blocks were split into windows by setting the parameter return_dataset = TRUE in the block_windowdataset() function. The haplotype variants within a given chromosome interval were recorded as 0, 1, 2 or 3 depending on the total number of haplotype variants present within the interval. In both cases, a mixed linear model implemented in R package GAPIT was used to detect the associations. To control for population structure in SNP-based analyses, 100,000 randomly selected markers were used to calculate the principal components. In haplotype-block-based GWAS, all haplotype blocks were used to calculate the principal components. In both cases, only the first three principal components were used to control the population structure. Two threshold options were used to identify significant associations including a more stringent Bonferroni correction and a less stringent Benjamini and Hochberg (1995) false discovery rate (FDR) at 5% significance level. To control for the effect of treatment and year, GWAS based on BLUPs was also performed and significant associations were reported only when there were SNP-trait or haplotype-trait association in at least two independent trials or in the BLUP-based analysis.

### 2.8 SNP-trait and haplotype-trait effects

Haplotype variation at loci with significant SNPs and their effects on traits in the introgression population were analyzed. The R package HaploBlocker v1.5.2 (Pook et al., 2019) was used to infer haplotypes at the genomic loci with significant SNP-trait associations. Heatmap.2 function provided in R package gplots was used to visualize the variation in haplotypes from hexaploid wheat and *Ae. tauschii*. However, at the *Ppd-D1* locus, visual comparison of the SNPs from the parental lines was done and the SNPs were annotated using snpEff v4.3 software to resolve haplotype variants in the *Ae. tauschii* lines that could not be clearly distinguished by HaploBlocker. SNPs significant at FDR ≤ 0.05 and their estimated allelic effects were selected from the association mapping results and used to verify if the haplotype effect corresponded to the observed phenotype in the introgression population. The mean and the standard deviation of the phenotype were calculated for each group of lines carrying a similar haplotype and the difference between the means was compared using Tukey’s honestly significant difference and the student’s *t*-test.

## 3 Results

### 3.1 Trait variation in the introgression population

Broad sense heritability (H^2^) of GY was 0.7 in 2018 and 0.64 in 2019, while for the yield component traits such as TGW, GA, GW and GL, H^2^ values were 0.85, 0.89, 0.83 and 0.95, respectively. The agronomic performance of the introgression lines relative to the controls (parents/checks) was assessed by comparing their yield and yield component traits under different treatments. The effect of treatment on yield was significant in 2018 (*P* < 2.2e-16), but not in 2019 (*P* = 0.24) at 95% confidence level. The latter is partially associated with more abundant rainfall in 2019 that reduced the difference in the water availability stress levels between the irrigated and non-irrigated field trials in Colby, KS. Based on the least squares means, significant differences in yield between controls and introgression lines was observed in 2019 and 2020, but not in 2018 (Table 1). Yield data collected from irrigated and rainfed (non-irrigated) field trials conducted between 2018 and 2020 revealed that up to 23% of the lines with introgressions produce more grain than the controls (Figure S2). In 2018 however, 3.2% and 48% of the introgression lines produced more grain than the parental lines and checks, respectively, in the non-irrigated trial (CO18), suggesting that the check cultivars were more sensitive to drought stress than the parental lines.

**Table 1.** Comparison of grain yield between introgression lines (progeny) and controls (checks/hexaploid parents) per treatment within a year using least squares (LS) means.

| Year | Treatment | Group | LSmean | SE | df | lower.CL | upper.CL | group |
| --- | --- | --- | --- | --- | --- | --- | --- | --- |
| 2018 | Irrigated | Progeny | 68.6 | 0.77 | 410 | 66.8 | 70.5 | a |
| 2018 | Irrigated | Check | 72.1 | 7.64 | 410 | 53.8 | 90.4 | a |
| 2018 | Irrigated | Parent | 75.7 | 3.6 | 410 | 67.1 | 84.3 | a |
| 2018 | Non-irrigated | Progeny | 50.1 | 6.43 | 395 | 34.6 | 65.5 | a |
| 2018 | Non-irrigated | Check | 51.6 | 0.66 | 395 | 50 | 53.2 | a |
| 2018 | Non-irrigated | Parent | 57.1 | 2.95 | 395 | 50 | 64.2 | a |
| 2019 | Irrigated | Progeny | 103 | 0.82 | 379 | 100.6 | 105 | a |
| 2019 | Irrigated | Parent | 119 | 3.8 | 379 | 109.5 | 128 | b |
| 2019 | Irrigated | Check | 121 | 9.05 | 379 | 99.3 | 143 | ab |
| 2019 | Non-irrigated | Progeny | 101 | 0.86 | 382 | 99.2 | 103 | a |
| 2019 | Non-irrigated | Parent | 116 | 3.85 | 382 | 106.3 | 125 | b |
| 2019 | Non-irrigated | Check | 116 | 9.44 | 382 | 93.5 | 139 | ab |
| 2020 | Non-irrigated | Progeny | 47.1 | 0.43 | 385 | 46 | 48.1 | a |
| 2020 | Non-irrigated | Parent | 55.3 | 2.01 | 385 | 50.5 | 60.1 | b |
| 2020 | Non-irrigated | Check | 59.3 | 4.79 | 385 | 47.8 | 70.8 | b |

**Note:** The 2018 and 2019 trials were conducted at Colby while the 2020 trial was conducted at Ashland, Kansas State. Groups with similar letters are not significantly different at 95% confidence level.

The proportion of introgression lines outperforming the checks and parental lines for the measured traits varied between treatments and years with a minimum of 0.8% for BM in the non-irrigated trial in 2019 (CO19) and a maximum of 73% for TGW in 2018 irrigated trial (COI18). The percentage increase in yield for the introgression lines that outperformed both checks and parents varied from 11% to 57%, while the number of lines that produced more grains varied from 6 to 94 (Table 2). Under drought stress conditions in 2018 (CO18), the mean yield of top performing introgression lines was 1.6 and 1.4-folds higher than the checks and parents, respectively (Figure 1). These results suggest that some introgression lines carry alleles that confer drought tolerance thus ensuring high productivity under stressful conditions. The highest yield potential of both introgression lines and controls was observed in 2019.The mean of the top yielding introgression lines reached 134 bushels per acre (BPA) while that of the parental lines and checks reached 119 and 121 BPA, respectively (Table 2).

**Table 2.**
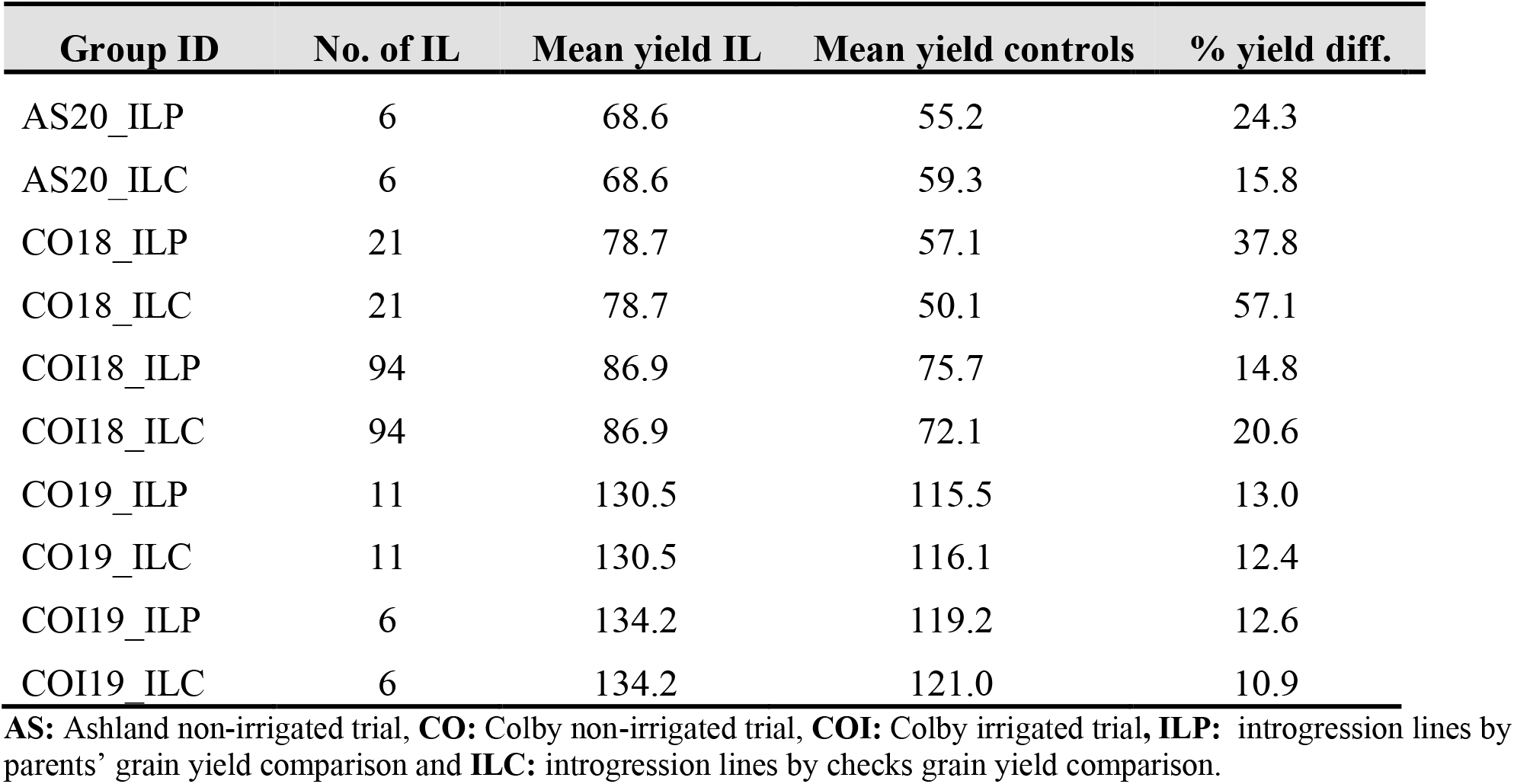
Percentage mean yield difference between top performing introgression lines and controls (parents and checks) per treatment in each year and location.

**Figure 1.**
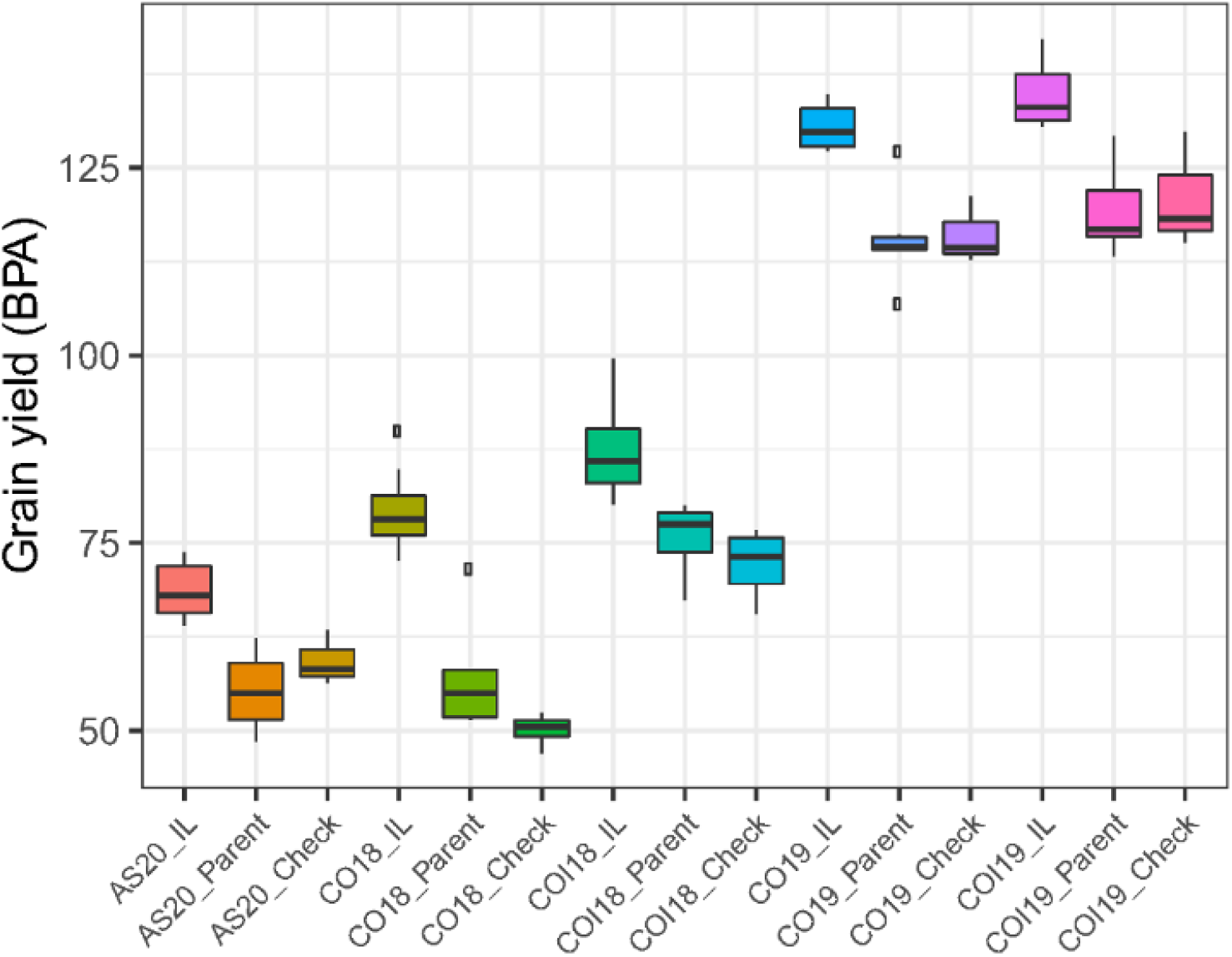
Boxplots comparing the mean grain yield between the top performing introgression lines (IL) and the controls (parents and checks) across treatments, years and locations. Where **AS20** refers to Ashland rainfed trial in 2020, **CO18** is Colby rainfed trial in 2018, **COI18** is Colby irrigated trial in 2018, **CO19** is Colby rainfed trial in 2019 and **COI19** is Colby irrigated trial in 2019.

The yield stability analyses were performed to identify introgression lines that are both high yielding and stable under various environmental conditions. Ranking by mean yield showed that 6% of the lines carrying introgression produced more grains than most parental lines, except Larry (File S1). The yield of these lines ranged from 84.4 to 92.7 BPA. The average rank (AR) of 16 stability statistics revealed 12 lines with introgressions showing high yield stability. Five of these lines were both stable and high yielding according to Kang’s rank-sum when compared to the controls. The yield of these five introgression lines (KS15SGDCB110-11, KS15SGDCB098-1, KS15SGDCB103-6, KS15SGDCB098-14 and KS15SGDCB111-1) varied between 82 and 93 BPA. The yield of the most stable and high yielding introgression line (92.7 BPA) was 9.8% higher than the average yield of the controls (84.4 BPA). These results indicate that novel alleles from *Ae. tauschii* have the potential to increase the adaptive potential of hard red winter wheat to different environmental conditions. In addition, the stability statistics could help to prioritize introgression lines for deployment in different agroecological zones depending on their ranking in stability and yield. Lines that are moderately high yielding but show good yield stability could be deployed in marginal environments, whereas less stable but high yielding lines could be deployed in less stressful environments to achieve high productivity.

Harvest index (HI), a measure of source-sink capacity was also assessed for stability in irrigated and rainfed trials. Ninety-two introgression lines showed a higher average HI (47.4-52.8) than the best parental line KS061406LN-26 (47.3). The average rank based on the 16 stability statistics placed 11 out of 92 lines in the top 20 most stable lines (File S2). Line KS15SGDCB111-1, which is high yielding and stable also ranked in the top five lines with stable and high average HI.

### 3.2 Trade-off between yield and yield component traits

Pearson’s correlation coefficients between average yield and yield stability based on average rank (AR) of the 16 stability statistics was -0.44 (*P* < 0.001), (File S1). However, the correlation between yield and Kang’s rank-sum (KR) was -0.71 (*P* < 0.001), indicating that the most stable introgression lines were not necessarily the highest grain yielders, although there were some exceptions. Similarly, the correlation between average HI and AR was -0.42 (*P* < 2.2e-16) while between HI and KR was -0.73 (*P* < 2.2e-16), (File S2).

The trade-off between yield and yield components was influenced by the treatment, year and location as evidenced by the variation in the levels of correlations (File S3). Higher positive correlations were observed among grain morphometric traits such as TGW, GA, GW and GL, ranging from 0.13 (between GW and GL) to 0.96 (between TGW and GA), (Figure 2). HI and GSW positively correlated with GY, while the correlation between GY and GN was positive but non-significant in all trials, except for Colby irrigated trial in 2019 (COI19) (File S3). BM correlated negatively with HI, but showed a positive correlation with GSW (File S3). In some cases, increase in the SNS resulted in a reduction in the TGW, GA, GW or GL, consistent with the previously observed trade-off between these traits (Kuzay et al., 2019). In contrast, HD positively correlated with the SNS and PH, which is in agreement with the previous findings (Shaw et al., 2013; Muqaddasi et al., 2019).

**Figure 2.**
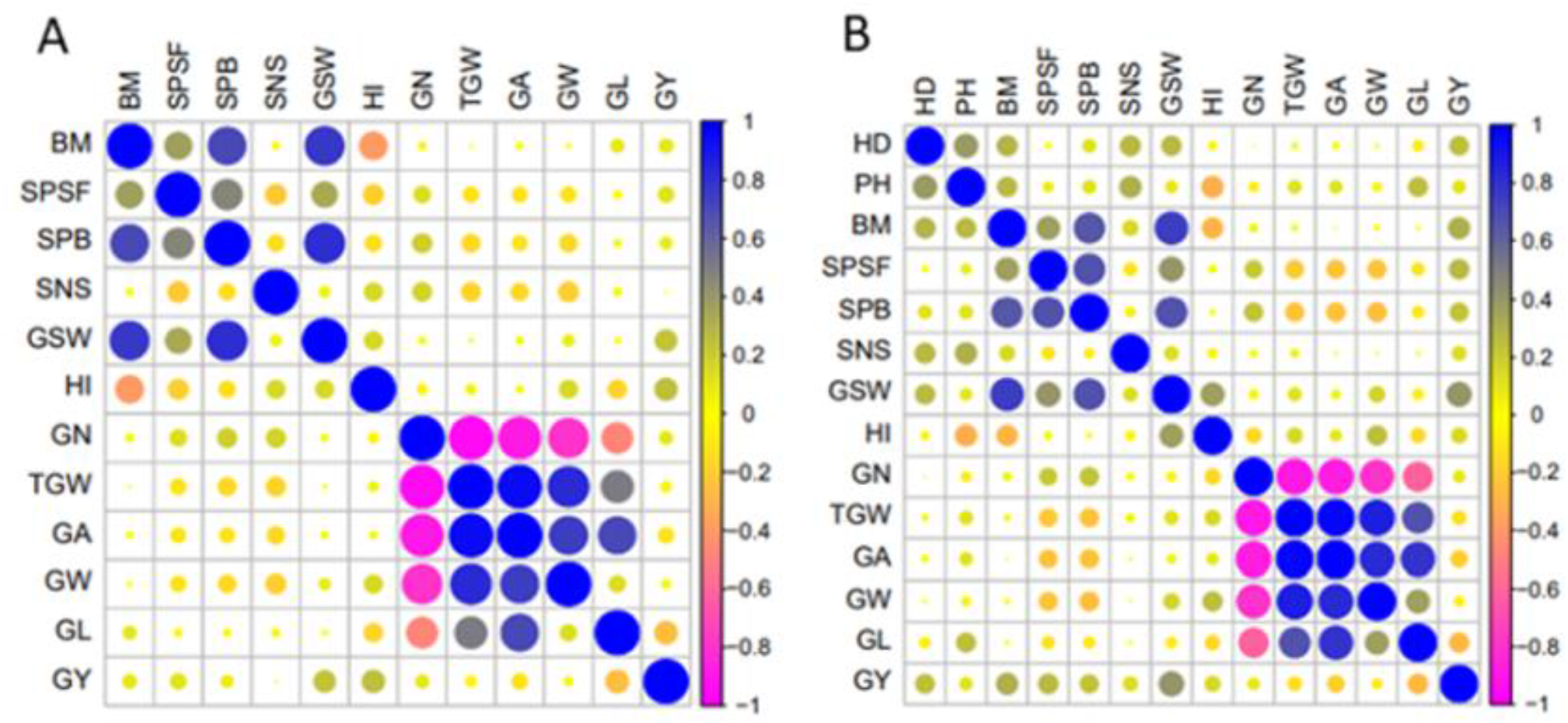
Pearson’s correlation coefficients between yield and yield component traits at Colby in 2018 rainfed trial (**A**) and Ashland in 2020 rainfed trial (**B**). Where **HD** is heading date, **PH** is plant height, **BM** is aboveground dry biomass, **SPSF** is spikes per square foot, **SPB** is spikes per bag, **SNS** is spikelet number per spike, **GSW** is grain sample weight, **HI** is harvest index, **GN** is grain number, **TGW** is thousand grain weight, **GA** is grain area, **GW** is grain width, **GL** is grain length, **GY** is grain yield.

To further understand the contribution of different yield components to the final yield, we compared the phenotypes of the top yielding introgression lines to those of the controls across all treatments (File S4). In the CO18 trial under non-irrigated conditions, the introgression lines that outperformed the controls in yield had the highest TGW, GA and GL, whereas under irrigated conditions (COI18), all yield component traits showed the highest levels of expression in the top yielding introgression lines. The top yielding introgression lines in the CO19 trial had the highest HI, GW, SNS and BM, while TGW and GA were comparable to those of the parental lines. In the COI19 trial, the TGW, GA and GW traits contributed more towards the final yield compared to the GL, HI and BM traits. In the AS20 trial, high levels of heterogeneity were observed among the top yielding lines for the TGW, GA, GW and GL traits. However, these lines showed higher BM than the controls, resulting in a reduced HI.

Previously, it was suggested that introgression from wild relatives might have negative impact on agronomic traits due to negative epistasis between the alleles of wild and cultivated wheat (Nyine et al., 2020). We investigated the relationship between the total size of introgressed genomic segments and phenotype. We found a positive linear relationship between GA, GL, SNS and the total size of the introgressed segments (Figure 3, File S5). For the TGW however, a positive linear relationship was only observed under drought stress conditions indicating that some wheat lines with large introgressions are efficient in utilizing the limited soil moisture and nutrients during grain filling. There was a negative relationship between GY, HI, GW, TGW under irrigated conditions and the size of introgression.

**Figure 3.**
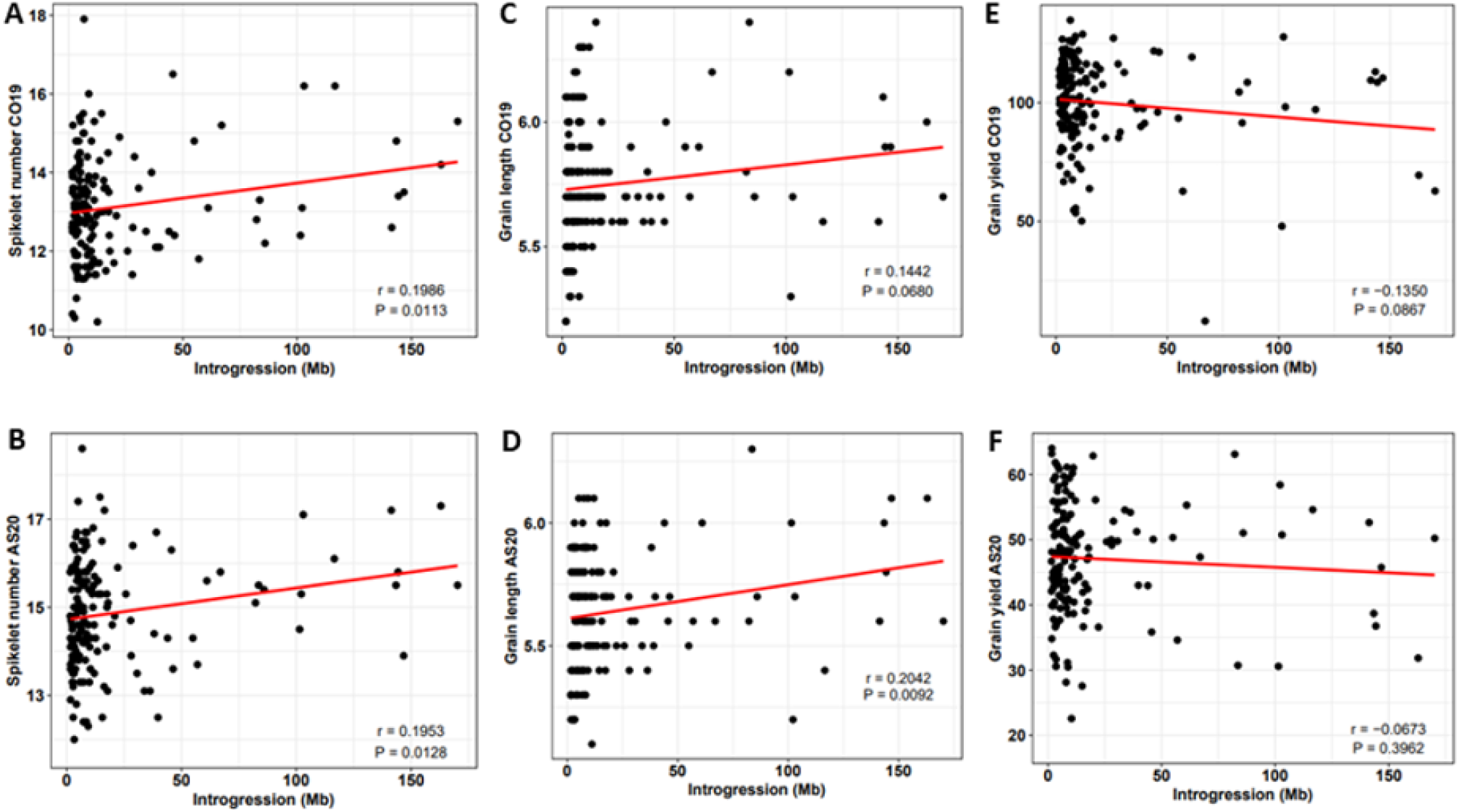
Relationship between spikelet number per spike (**A & B**), grain length (**C & D**), grain yield (**E & F**) and the proportion of introgression under non-irrigated conditions at Colby in 2019 (**CO19**) and Ashland in 2020 (**AS20**). Here, **r** is the correlation coefficient and **P** is the significance of the correlation between introgression size and observed trait phenotype.

### 3.3 Genotyping and imputation

To identify *Ae. tauschii* haplotypes in the D genome of introgression population, we generated high-density SNP data. By whole-genome sequencing of six hexaploid parental lines and 21 *Ae. tauschii* accessions used for generating octoploid parents, we identified about 20 million high-quality SNP variants (MAF ≥ 0.05) and used them for genotype imputation in the introgression population genotyped by complexity-reduced sequencing. The total number of D genome SNPs retained after filtering out SNPs with genotype probability below 0.7 and MAF < 0.05 was 5.2 million.

### 3.4 Haplotypic variation between ssp. *strangulata* and ssp. *tauschii* families

Using HaploBlocker v1.5.2, we identified 4,764 and 6,429 non-overlapping haplotype blocks in the *Ae. tauschii* ssp. *strangulata* (FAM93) and *Ae. tauschii* ssp. *tauschii* (FAM97) families, respectively. After filtering out the monomorphic haplotypes between the parental lines, 869 (18%) and 3,020 (47%) segregating haplotypes were retained in FAM93 and FAM97, respectively (Table 3, File S6). The low proportion of segregating haplotypes between hexaploid wheat and ssp. *strangulata* D genomes is in agreement with the finding that *Ae. tauschii* ssp. *strangulata* was the donor of the D genome of hexaploid wheat (Wang et al., 2013). These results also suggest that the high similarity between the genome of ssp. *strangulata* and the D genome of hexaploid wheat could result in underestimation of the proportion of introgressed haplotypes. The average genome-wide haplotype block length in FAM93 was higher (2 Mb) than that in the FAM97 (1 Mb), (File S6). There was a significant difference in the introgressed haplotype length between lines in FAM93 and FAM97 based on the t-test (*P* = 3.1e-16). The longest haplotype introgressed in all lines from FAM93 was 44 Mb on chromosome arm 3DL while in FAM97 only four lines had a haplotype >32 Mb on chromosome arm 7DL. The number of segregating haplotypes in FAM93 varies from 32 (3D) to 336 (2D), while in FAM97 the number of segregating haplotypes varies from 173 (3D) to 617 (5D), (Table 3). In FAM93 and FAM97, the average frequency of each haplotype from *Ae. tauschii* parents in the introgression lines was 11 and 4, respectively (File S6).

**Table 3.** Variation of introgressed haplotypes between *Ae. tauschii* ssp. *strangulata* (FAM93) and *Ae. tauschii* ssp. *tauschii* (FAM97) families.

| Family | Pedigree | Subspecies | Chrom | NH | MHL (bp) | MHF | Range (bp) |
| --- | --- | --- | --- | --- | --- | --- | --- |
| FAM93 | KanMark x TA1642 | *strangulata* | chr1D | 105 | 1,896,815 | 4.6 | 1,788-8,639,976 |
| FAM93 | KanMark x TA1642 | *strangulata* | chr2D | 336 | 1,326,620 | 20.0 | 813-30,656,640 |
| FAM93 | KanMark x TA1642 | *strangulata* | chr3D | 32 | 6,653,571 | 4.9 | 2,525-44,264,371 |
| FAM93 | KanMark x TA1642 | *strangulata* | chr4D | 143 | 1,865,609 | 4.3 | 2,942-12,070,089 |
| FAM93 | KanMark x TA1642 | *strangulata* | chr5D | 138 | 2,302,767 | 7.1 | 556-15,103,840 |
| FAM93 | KanMark x TA1642 | *strangulata* | chr6D | 65 | 2,519,945 | 3.0 | 3,455-13,483,300 |
| FAM93 | KanMark x TA1642 | *strangulata* | chr7D | 50 | 2,667,416 | 2.04 | 1,627-11,349,447 |
| FAM97 | Danby x TA2388 | *tauschii* | chr1D | 495 | 829,609 | 3.1 | 762-9,169,049 |
| FAM97 | Danby x TA2388 | *tauschii* | chr2D | 352 | 1,305,439 | 1.8 | 813-15,991,790 |
| FAM97 | Danby x TA2388 | *tauschii* | chr3D | 173 | 2,968,356 | 3.0 | 884-27,225,468 |
| FAM97 | Danby x TA2388 | *tauschii* | chr4D | 540 | 750,352 | 0.8 | 1,982-20,010,593 |
| FAM97 | Danby x TA2388 | *tauschii* | chr5D | 617 | 770,483 | 10.1 | 1,243-15,396,133 |
| FAM97 | Danby x TA2388 | *tauschii* | chr6D | 373 | 830,168 | 3.8 | 764-23,208,199 |
| FAM97 | Danby x TA2388 | *tauschii* | chr7D | 465 | 1,000,899 | 3.7 | 1,397-32,547,936 |

**NH** = number of haplotype blocks, **MHL** = mean haplotype block length, **MHF** = mean haplotype frequency

### 3.5 SNP- and haplotype-based genome-wide association mapping

Genome-wide association study was performed in the *Ae. tauschii* introgression population to assess the effects of introgression into the D genome on the variance of traits related to biomass, yield and yield components and tenacious glume. The marker-trait association analyses were based on both individual SNPs and haplotype blocks identified by HaploBlocker from the 5.2 million imputed variants. We report only those associations that are replicated in at least two independent field trials and show significant association with both SNPs and haplotypes at FDR 0.05 (Table S1). Several genomic loci with significant associations distributed on the D genome chromosomes were detected for GL, GW and SNS. For other traits such as GY, TGW, GN, GA, HI, BM, GSW and SPSF, no consistent associations replicated in independent trials were detected.

We identified multiple significant SNP-trait and haplotype-trait associations from all trials on chromosome arms 2DS and 7DS for GL (Figure 4). The most significant SNPs were located in haplotype block windows 22,262,355 – 22,289,017 bp, 30,582,113 – 30,595,115 bp and 80,864,297 – 81,398,316 bp on chromosome arm 2DS, and 11,024,311 – 11,374,767 bp on chromosome arm 7DS (Table S1). Association analysis based on BLUPs confirmed the results from individual trials for GL on these two chromosomes. Other significant associations detected in at least two trials were identified on chromosome arms 3DS and 5DS (File S7).

**Figure 4.**
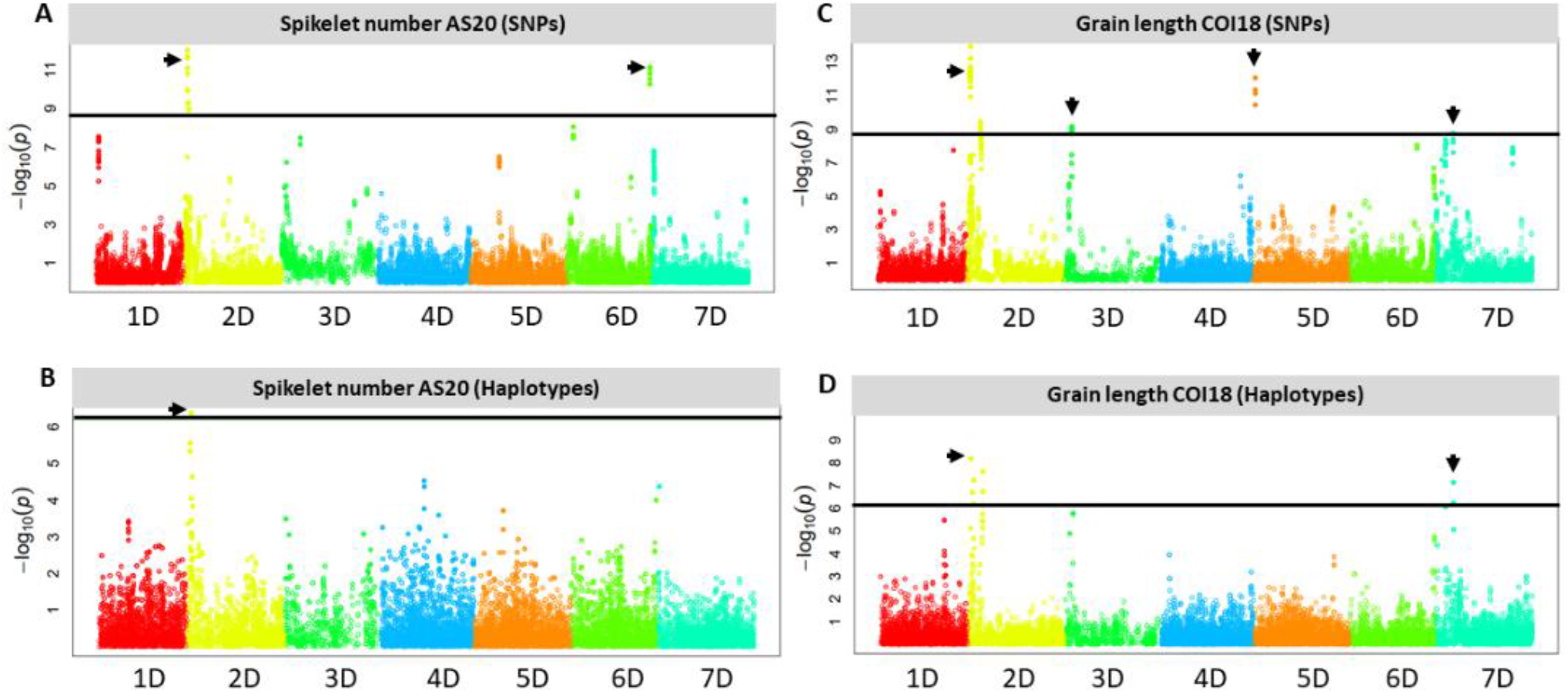
Manhattan plots showing the D genome loci with SNPs and haplotypes that are significantly associated with spikelet number per spike at Ashland in 2020 (**AS20**) under rainfed conditions (**A & B**) and grain length at Colby in 2018 (**COI18**) under irrigated conditions (**C & D**) in the *Ae. tauschii* introgression population. The horizontal solid black line shows a threshold of 0.05 significance level for Bonferroni correction, the black arrowheads indicate the SNPs and haplotypes above the threshold.

We identified haplotypes with significant SNPs associated with GW on 1DL, 2DS, 6DL and 7DS from at least two independent trials that were confirmed by BLUP-based analysis (File S7).

Haplotype block windows 65,964,778 – 66,124,103 bp and 66,265,325 – 66,266,089 bp showed the most significant association on 2DS.

At 95% confidence level, the most significant SNP-trait associations were identified on chromosome arms 2DS and 6DL for SNS from three independent trials (COI19, CO19 and AS20), (Figure 4, File S7). The most significant associations are located at 16.5 Mb and 463.8 Mb on 2DS and 6DL, respectively. Haplotype-trait analysis confirmed the association on 2DS for SNS at the 16.5 Mb locus located within the haplotype block window 16,497,666 – 16,548,006 bp. At FDR < 0.05, there was no haplotype block window on 6DL locus that overlapped with the significant SNP-trait association.

Previous studies have shown that SNS is linked to HD (Shaw et al., 2013; Muqaddasi et al., 2019). In the current study, we detected significant associations with SNS on chromosome arms 2DS and 6DL. We had one year data for HD and PH collected from Ashland in 2020, which provided us a good opportunity to validate this link in the *Ae. tauschii* introgression population. Genome-wide association mapping detected significant associations with HD on chromosome arms 2DS and 4DL while all D genome chromosomes showed significant association with PH but the strongest signals were observed on 1DS, 3DS and 6DL. The haplotype block window 16,548,753 – 16,639,561 bp on 2DS with the most significant SNPs for HD overlapped the locus showing significant association with SNS, which is in close proximity to another haplotype block overlapping with the most significant SNPs for SNS (16,497,666 – 16,548,006 bp). These results suggest that the expression of these two traits could be co-regulated.

For HD, the haplotype block windows on chromosome arm 4DL 442,735,095 – 442,751,954 bp and 459,271,685 – 459,290,731 bp had the most significant SNP-trait associations. The three traits (SNS, HD and PH) are known to be affected by the *Rht8* and *Ppd-D1* genes on 2DS, in addition to *Rht1* on 4D, which controls plant height and flowering time (Borojevic and Borojevic 2005; Chen et al., 2018). Due to the lack of SNPs located near the *Ppd-D1* gene locus at ∼34Mb (33,961,438 – 33,951,651 bp interval in CS RefSeq v1.0), we could not directly validate its association with these traits. However, significant associations for SNS were detected at ∼3 Mb next to the *Ppd-D1* locus in the CO19 and AS20 trials on haplotype blocks 2D:30,192,335 – 30,264,745 bp and 2D:28,829,778 – 28,937,705 bp, respectively. In the parental lines with high density SNPs (∼20 million), the *Ppd-D1* locus had SNPs, which allowed us to precisely map the haplotypes from *Ae. tauschii* and hexaploid wheat lines. Results from HaploBlocker showed that all hexaploid parents carry an identical haplotype, which is distinct from that of *Ae. tauschii* accessions.

Using SNPs identified by whole-genome sequencing of parental lines, we characterized haplotypic diversity at the *Ppd-D1* locus (Figure 5A). All hexaploid wheat lines carried the same *Ppd-D1* haplotype (Hap1), while seven haplotypes of the *Ppd-D1* gene (Hap2 -Hap8) were identified in *Ae. tauschii*. Whole genome sequencing of 21 *Ae. tauschii* revealed broader range of *Ppd-D1* diversity compared to a previous study (Guo et al., 2009), which identified only three *Ppd-D1* haplotypes. The *Ae. tauschii* ssp. *strangulata* accessions carried haplotypes that were identical to hexaploid wheat, except for Hap2 in TA1642, which had one SNP at position 33,952,131 bp (Figure 5A). The *Ppd-D1* genic region in *Ae. tauschii* ssp. *tauschii* accessions has one synonymous (SN), three intronic (IN) and one missense (MS) SNPs. The missense variant at position 33,955,614 bp results in His16Asn change, which is predicted to have moderate functional impact, and only present in lines with haplotype Hap5 (Figure 5A). Next, we inferred the parental haplotypes of the *Ppd-D1* locus in the introgression population by using SNPs within the ∼1-2 Mb region surrounding the *Ppd-D1* locus. About 82% of the introgression lines carried haplotype Hap1.

**Figure 5.**
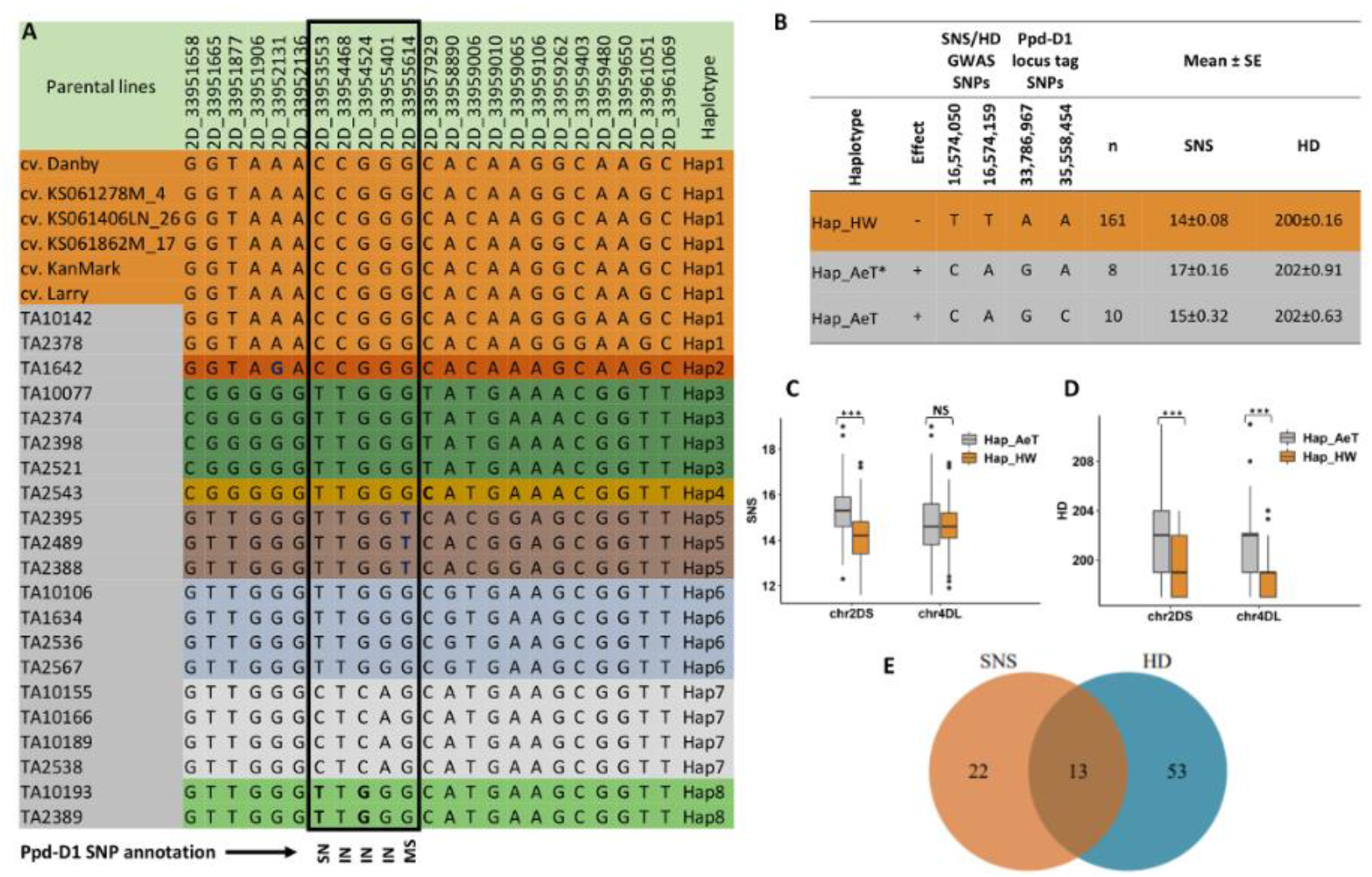
Effect of haplotypes introgressed from *Ae. tauschii* into the chromosome arms 2DS and 4DL of hexaploid wheat on the spikelet number per spike (SNS) and heading date (HD) of the introgression lines and the possible link to *Ppd-D1* gene located on 2DS. **(A)** SNP table showing the haplotype variants at the *Ppd-D1* gene locus in winter wheat accessions (top six) and the 21 *Ae. tauschii* lines. The black rectangle shows SNPs within the coding region of the gene where **SN** is a synonymous SNP, **IN** is an intronic SNP and **MS** is a missense SNP (His16Asn) as reported by snpEff v4.3 software. **(B)** Unique haplotypes in the introgression population tagging the *Ppd-D1* locus and GWAS signal for SNS and HD on chromosome arm 2D, and the associated phenotype. Hap_HW is present in introgression lines that have Hap1 from hexaploid wheat at the *Ppd-D1* locus, Hap_AeT includes lines that have Hap3-6 and 8 while Hap_AeT* includes lines that have Hap2 and Hap7 at the *Ppd-D1* locus in *Ae. tauschii* parents. The TT and CA alleles at the GWAS signal have reducing and increasing effects, respectively, on SNS and HD. **(C)** Boxplot showing the impact of introgression from *Ae. tauschii* in chromosome arms 2DS and 4DL on SNS. **(D)** Boxplot showing the impact of introgression from *Ae. tauschii* in chromosome arms 2DS and 4DL on HD. **(E)** A Venn diagram showing the number of introgression lines in the 90th percentile for SNS and HD. Lines in the intersection have the increasing alleles on both 2DS and 4DL loci associated with SNS and HD. *** indicates significant difference between groups with P < 0.001 while NS indicates a nonsignificant difference based on t-test statistics.

Further, we evaluated linkage of *Ppd-D1* haplotypes with SNP alleles showing significant association with variation in SNS and HD. For this purpose, we used two SNP sites, 2D_33786967 and 2D_35558454, that flank the *Ppd-D1* locus on both sides and have genotyping information in the introgression population. We compared them to SNP alleles that were significantly associated with SNS and HD in haplotype block window 2D: 16,548,753 – 16,639,561 bp (2D_16574050 and 2D_16574159), spanning ∼17 Mb region (Figure 5B). We found that the GWAS alleles associated with increase in SNS and HD in the introgression population are also linked with two *Ae. tauschii* haplotypes (Hap_AeT* and Hap_AeT), whereas the GWAS alleles associated with decreasing effects were in LD with Hap_HW contributed by the hexaploid wheat parents. The Hap_AeT* group of haplotypes was contributed by the *Ae. tauschii* parents having Hap2 and Hap7 at the *Ppd-D1* locus.

### 3.6 The phenotypic effects of haplotype block variants

#### 3.6.1 Average spikelet number per spike (SNS) and Heading date (HD)

Significant haplotype-trait associations were identified on chromosome arms 2DS and 4DL that influence SNS and HD. Chromosome 2DS had multiple introgressed haplotypes that are significantly associated with the variation in SNS and HD with the most significant haplotypes located at 16,497,666 – 16,548,006 bp and 16,548,753 – 16,639,561 bp for SNS and HD, respectively. The haplotype variants with the increasing effect at these loci were from *Ae. tauschii* parents while those with reducing effect were from the hexaploid wheat lines (Figure 5C, 5D). The verification of GWAS results for allelic effect at 2DS locus associated with SNS and HD supports the above observation (File S8). We observed a positive Pearson’s correlation coefficient between SNS and HD; and lines having haplotypes from either parent showed significant differences in the phenotype based on a t-test (*r* = 0.23, *P* = 3.31e-07) at 95% confidence level. Haplotypes on 4DL had smaller effect on SNS compared to HD. Among 35 and 66 introgression lines having SNS and HD trait values above the 90^th^ percentile of trait distribution, respectively, 13 lines had the increasing alleles from *Ae. tauschii* at both 2DS and 4DL loci associated with SNS and HD traits (Figure 5E).

Initially, we did not detect a significant GWAS signal directly associated with SNPs within the *Ppd-D1* gene due to the lack of high-quality imputed SNPs in this region. Further analysis of parental haplotypes identified SNP variants linked with both the *Ppd-D1* haplotypes and haplotypes at 28 Mb and 30 Mb region showing the significant haplotype-trait association in two trials. These haplotypes were within ∼3 Mb from the *Ppd-D1* locus and likely overlap with *Ppd-D1*. We performed the analysis of variance to determine the effect of different haplotype variants identified in the parental accessions on the SNS in the introgression population using data from three experimental trials (COI19, CO19 and AS20) (Table 4). Results show that both hexaploid and *Ae. tauschii* haplotypes have a significant effect on SNS (*P* < 0.001) (Table 4, Figure 5B). Among the *Ae. tauschii* haplotypes, Hap7 had the highest impact on SNS (*P* = 0.003) followed by Hap2 (*P* = 0041) and Hap3 (*P* = 0.040). In contrast, Hap5 with the His16Asn missense mutation had a negative effect on SNS and was not significantly different from Hap1 present in hexaploid wheat lines (*P* = 0.072). These results are consistent with earlier studies that showed that the *Ppd-D1* gene located at ∼34 Mb (33,961,438 – 33,951,651 bp interval in CS RefSeq v.1.0), that plays a role in flowering time regulation in wheat also has a strong effect on variation in the SNS (Beales et al., 2007, Guo et al., 2009).

**Table 4.**
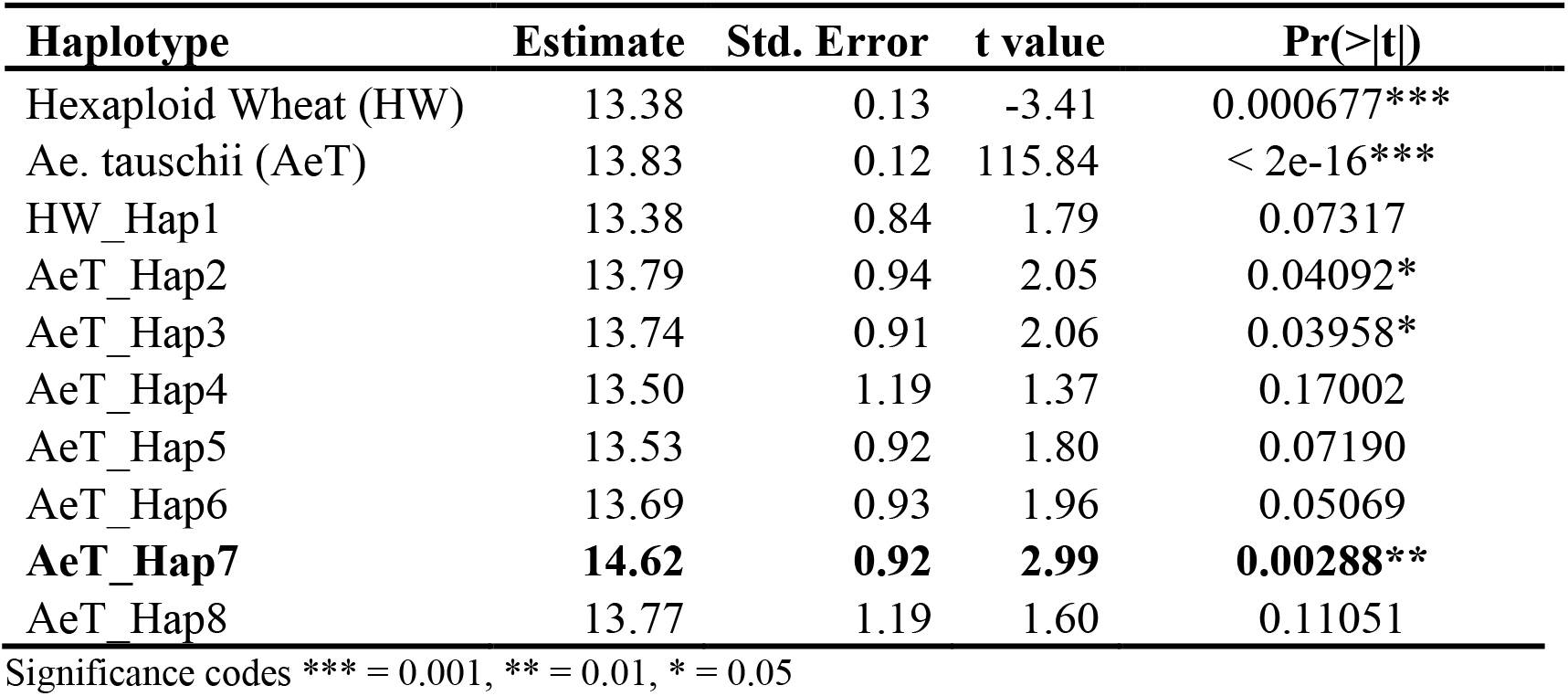
Analysis of variance for the effect of *Ppd-D1* haplotype variants from hexaploid wheat and *Ae. tauschii* on the spikelet number per spike in three experimental trials of the introgression population.

| Haplotype | Estimate | Std. Error | t value | Pr(>\|t\|) |
| --- | --- | --- | --- | --- |
| Hexaploid Wheat (HW) | 13.38 | 0.13 | -3.41 | 0.000677*** |
| *Ae. tauschii* (AeT) | 13.83 | 0.12 | 115.84 | < 2e-16*** |
| HW_Hap1 | 13.38 | 0.84 | 1.79 | 0.07317 |
| AeT_Hap2 | 13.79 | 0.94 | 2.05 | 0.04092* |
| AeT_Hap3 | 13.74 | 0.91 | 2.06 | 0.03958* |
| AeT_Hap4 | 13.50 | 1.19 | 1.37 | 0.17002 |
| AeT_Hap5 | 13.53 | 0.92 | 1.80 | 0.07190 |
| AeT_Hap6 | 13.69 | 0.93 | 1.96 | 0.05069 |
| **AeT_Hap7** | **14.62** | **0.92** | **2.99** | **0.00288\*\*** |
| AeT_Hap8 | 13.77 | 1.19 | 1.60 | 0.11051 |

Significance codes \*\*\* = 0.001, \*\* = 0.01, \* = 0.05

#### 3.6.2 Pleiotropic effects of haplotypes on yield component traits

We also evaluated the effects of distinct haplotypes associated with HD on other traits. Haplotype Hap_AeT from chromosome 2D located at 16,548,753 – 16,639,561 bp is associated with significant increase in the days to heading and SNS without significant impact on BM, HI and GSW (Table 5).

**Table 5.** Chromosome 2D haplotypes variants associated with the spikelet number and heading date and how they influence other traits in the introgression population.

| Trait | Haplotype | Loc1 Freq. | chr2D:16548753-16639561 | SD |
| --- | --- | --- | --- | --- |
| HD | Hap_HW | 53 | 199<sup>b</sup> | 2.28* |
| **HD** | **Hap_AeT** | **156** | **202<sup>a</sup>** | **2.86↓** |
| HD | Hap_HW/AeT | 142 | 200<sup>c</sup> | 2.29‡ |
| PH | Hap_HW | 53 | 68.27<sup>ab</sup> | 6.79 |
| PH | Hap_AeT | 156 | 70.82<sup>a</sup> | 6.97 |
| PH | Hap_HW/AeT | 142 | 65.9<sup>b</sup> | 8.68 |
| BM | Hap_HW | 53 | 155.18<sup>a</sup> | 28.21 |
| BM | Hap_AeT | 156 | 156.26<sup>a</sup> | 31.01 |
| BM | Hap_HW/AeT | 142 | 157.52<sup>a</sup> | 35.82 |
| SNS | Hap_HW | 53 | 14.72<sup>b</sup> | 1.2 |
| **SNS** | **Hap_AeT** | **156** | **15.27<sup>a</sup>** | **1.12** |
| SNS | Hap_HW/AeT | 142 | 14.05<sup>c</sup> | 1.03 |
| GSW | Hap_HW | 53 | 66.84<sup>a</sup> | 14.71 |
| GSW | Hap_AeT | 156 | 65.82<sup>a</sup> | 13.67 |
| GSW | Hap_HW/AeT | 142 | 66.47<sup>a</sup> | 13.57 |
| HI | Hap_HW | 53 | 42.85<sup>a</sup> | 4.35 |
| HI | Hap_AeT | 156 | 42.23<sup>a</sup> | 4.24 |
| HI | Hap_HW/AeT | 142 | 42.89<sup>a</sup> | 6.28 |

\*Haplotype with reducing effect ↓Haplotype with increasing effect ‡Haplotype with intermediate effect, **Loc1** = chr2D:16,575,276 – 16,635,669 bp, **Loc1 Freq.** = number of introgression lines having the haplotype, **HD** is heading date, **PH** is average plant height, **BM** is aboveground total dry biomass, **SNS** is spikelet number per spike, **GSW** is Grain sample weight from biomass sample and **HI** is harvest index. Means with the same superscript letters are not statistically significant for each trait based on Tukey's honestly significant difference at 95% confidence level.

We compared the effects of two haplotypes associated with GW on 2DS (2D: 65,964,778 – 66,124,103 bp and 2D: 66,265,325 – 66,266,089 bp), where Hap_HW from hexaploid parents and Hap_AeT_st from *Ae. tauschii* ssp. *strangulata* increase GW and Hap_AeT_ta from the *Ae. tauschii* ssp. *tauschii* parents reduces GW (Table 6). In Colby 2018 non-irrigated trial data, the haplotype at the 2D: 65,964,778 – 66,124,103 bp locus that was associated with increase in GW and GL was also linked with increase in grain area, and decrease in grain number. While both Hap_AeT_st and Hap_AeT_ta haplotypes at 2D: 65,964,778 – 66,124,103 bp were associated with increase in grain length, only Hap_AeT_ta was linked with the significant increase in grain number (Table 6). The Hap_AeT_ta haplotype at the 2D: 66,265,325 – 66,266,089 bp haplotype block had similar effects on GN, although the observed difference was not significant.

**Table 6.** Chromosome 2D haplotypes variants associated with the grain width in Colby 2018 rainfed trial and their effects on other traits in the introgression population.

| Trait | Haplotype | Haploblock chr2D:65964778–66124103 |  |  | Haploblock chr2D:66265325–66266089 |  |  |
| --- | --- | --- | --- | --- | --- | --- | --- |
|  |  | Freq. | Trait mean | Trait SD | Freq. | Trait mean | Trait SD |
| GN | Hap_AeT_st | 5 | 475.50<sup>b</sup> | 27.86 | 20 | 504.15<sup>a</sup> | 55.28 |
| GN | Hap HW | 140 | 499.04<sup>b</sup> | 44.3 | 304 | 502.97<sup>a</sup> | 41.6 |
| GN | Hap AeT_ta | 202 | 510.94<sup>a</sup> | 43.38 | 27 | 540.37<sup>a</sup> | 47.94 |
| TGW | Hap AeT_st | 5 | 32.74<sup>a</sup> | 2.15 | 20 | 31.54<sup>a</sup> | 2.74 |
| TGW | Hap HW | 140 | 31.40<sup>a</sup> | 2.2 | 304 | 31.30<sup>a</sup> | 2 |
| TGW | Hap AeT_ta | 202 | 31.01<sup>a</sup> | 1.98 | 27 | 29.54<sup>a</sup> | 1.76 |
| GA | Hap_AeT_st | 5 | 14.38<sup>a</sup> | 1.18 | 20 | 13.89<sup>a</sup> | 1.03 |
| GA | Hap HW | 140 | 13.50<sup>b</sup> | 0.69 | 304 | 13.53<sup>a</sup> | 0.66 |
| GA | Hap AeT_ta | 202 | 13.53<sup>b</sup> | 0.66 | 27 | 13.28<sup>a</sup> | 0.61 |
| GW | Hap AeT_st | 5 | 3.14<sup>a</sup> | 0.05 | 20 | 3.09<sup>ab</sup> | 0.08Δ |
| GW | Hap HW | 140 | 3.13<sup>a</sup> | 0.09 | 304 | 3.09<sup>a</sup> | 0.09Δ |
| GW | Hap AeT_ta | 202 | 3.06<sup>b</sup> | 0.08 | 27 | 3.00<sup>b</sup> | 0.07# |
| GL | Hap AeT_st | 5 | 6.25<sup>a</sup> | 0.51 | 20 | 6.11<sup>a</sup> | 0.4 |
| GL | Hap HW | 140 | 5.85<sup>b</sup> | 0.22 | 304 | 5.93<sup>b</sup> | 0.24 |
| GL | Hap AeT_ta | 202 | 6.00<sup>a</sup> | 0.24 | 27 | 6.01<sup>a</sup> | 0.22 |
| GY | Hap AeT_st | 5 | 59.19<sup>a</sup> | 14.46 | 20 | 58.76<sup>a</sup> | 14.77 |
| GY | Hap HW | 140 | 60.51<sup>a</sup> | 12.04 | 304 | 61.26<sup>a</sup> | 12.03 |
| GY | Hap_AeT_ta | 202 | 61.28<sup>a</sup> | 12.54 | 27 | 59.55<sup>a</sup> | 13.41 |

Δ haplotype that increases grain width, # haplotype that reduces grain width. Means with the same superscript letters are not statistically significant for each trait based on Tukey’ s honestly significant difference at 95% confidence level.

Similarly, haplotype block 6D: 463,775,852 – 463,809,722 bp associated with PH and SNS has three variants (File S7). Haplotype variant Hap1_HW&AeT is associated with increase in SNS, PH, BM and GY. It showed no association with HD and grain traits, except grain number where an intermediate effect was observed. This haplotype variant is present in two *Ae. tauschii* lines (TA2389 and TA2398) and the hexaploid parents excluding KanMark and KS061862M-17. Haplotype variant Hap0_HW&AeT, which is found in KanMark and KS061862M-17 and two ssp. *strangulata* accessions (TA1642 and TA2378) has an intermediate effect on GY, reducing it by 2 bushels compared to Hap1_HW&AeT. The third haplotype variant (Hap_AeT) is present only in the *Ae. tauschii* lines.

The haplotype Hap_AeT contributed by *Ae. tauschii* at haplotype block 7D:14,185.651 – 14,596,748 bp, was associated with significant increase in SNS compared to haplotypes present in winter wheat (Table S2). This increase was associated with significant decrease in GL and had no significant effect on GY. At 7D:14,722,457 – 14,817,138 bp, the Hap_AeT haplotype contributed by *Ae. tauschii* was also linked with significant increase in SNS compared to Hap1_HW&AeT detected in both hexaploid wheat and *Ae. tauschii* parents. However, increase in SNS for this haplotype was connected with decrease in both GL and GY. At this haplotype block (7D:14,722,457 – 14,817,138 bp), the most significant increase in GY was observed for lines carrying haplotype Hap0_HW&AeT, which was associated with moderate increase in both SNS and GL (Table S2).

## 4 Discussion

Here, we performed the sequence-based analysis of haplotypes in the wild-relative introgression population developed by crossing a diverse panel of *Ae. tausc*hii accessions with winter wheat cultivars. Our results demonstrate that combining whole genome sequencing of wild and cultivated wheat founder lines with the complexity-reduced sequencing of a derived introgression population provides an effective framework for SNP imputation. Because most breeding populations are based on a limited number of founders, often including 10-30 lines, their whole genome sequencing is feasible in crops even with such large genomes as wheat, and provides a comprehensive description of allelic diversity present in a breeding population. The latter makes sequenced founders an ideal reference panel for imputing genotypes in a breeding population genotyped using low-coverage or complexity-reduced sequencing. This was recently demonstrated by imputing genotypes in the wheat MAGIC population genotyped by low-coverage sequencing (Scott et al., 2021). The composition of our introgression population including multiple bi-parental cross families (Nyine et al., 2020) also shifts the population allele frequency towards more common variants, which could be imputed with a higher accuracy than rare variants (Huang et al., 2015). In addition, the high levels of LD in the introgression population should increase the length of haplotype blocks and facilitate detection of matching haplotypes in the reference panel of founders using even sparse genotyping data generated by low-coverage or complexity-reducing sequencing. Consistent with these assumptions, an imputation algorithm implemented in Beagle (Browning and Browning 2013) allowed us to impute nearly 5.2 million SNPs in the introgression population with high genotype call probabilities above 0.7 using SNPs generated by complexity-reduced sequencing of this population and nearly 20 million variants identified in 27 founders. This high-density SNP marker data permitted detailed characterization of introgressed haplotypes (Pook et al., 2019) and assessing their effects on productivity traits.

Our results demonstrate that wild relative introgressions into the D genome of wheat, the least diverse amongst the three subgenomes, (Chao et al., 2010; Singh et al., 2019; Jordan et al., 2015) is associated with the increased levels of variation in yield and yield component traits. Analysis of data from several years and locations under irrigated and non-irrigated conditions revealed many superior introgression lines that produce more grains or show higher yield stability than the control cultivars. The yield increase in top-performing introgression lines was driven by a combination of yield component traits and in many cases, it was associated with increased grain size, grain weight, and biomass or improved harvest index. These results suggest that wild-relative introgression has the potential to positively affect source-sink balance, which was suggested to be one of the important factors contributing to yield potential (Reynolds et al., 2017). Many of these high yielding lines (∼23%) were also among the top lines showing the highest levels of yield stability, indicating that introgression from *Ae. tauschii* likely improves the adaptive potential of hard red winter wheat in different environmental conditions. Consistent with this conclusion, the highest impact of introgression on yield was found in a non-irrigated trial, indicating that alleles from *Ae. tauschii* likely improve the adaption of hexaploid wheat to water-limiting conditions. The *Ae. tasuchii* accessions used to generate the introgression population represent both L1 and L2 lineages (Wang et al., 2013) and originate from a broad range of geographical locations with variable climatic conditions, likely capturing adaptive haplotypes from regions prone to drought stress.

Heading date is one of the key agronomic traits linked with wheat adaptation to different geographical locations and improvement in yield (Jung and Müller 2009). In our population, a positive correlation was observed between heading date and the spikelet number per spike with some lines showing up to two-week delay in heading date. Several haplotype blocks on chromosome arms 2DS and 4DL were significantly associated with variation in spikelet number and heading date. The haplotypes with increasing effects at both loci were derived from the *Ae. tauschii* indicating their potential for modulating both traits in bread wheat. Chromosome 2DS is known to harbor the *Ppd-D1* and *Rht8* genes that control flowering time and plant height, respectively, and also could affect spikelet number (Shaw et al., 2013; Muqaddasi et al., 2019). The overlapping haplotype blocks associated with spikelet number and heading date were identified on 2DS, confirming that the two traits co-segregating in the population have a common genetic basis. We demonstrated that these 2DS haplotypes are associated with different allelic variants of the *Ppd-D1* gene from *Ae. tauschii*. These results are consistent with the earlier studies that demonstrated that different alleles of the *Ppd-D1* gene have distinct effects on heading date and spikelet number per spike (Beales et al., 2007, Guo et al., 2009). These effects correlated with the relative expression levels of each *Ppd-D1* allele (Guo et al., 2009), suggesting that functional mutations within the *Ppd-D1* coding region and the modifier mutations in the regulatory region of the gene likely account for the variation in these traits in the *Ae. tauschii*-winter wheat introgression population. The developmental plasticity modulated by *Ppd-D1* is mediated by changes in the expression of flowering time genes (Gol et al. 2021). It was shown that the *Ppd-H1* from wild barley is capable of integrating environmental signals to control heading date and minimize the negative impact of transient drought stress on spikelet number (Gol et al. 2021). Consistent with this observation, in the current study, introgression lines that have a high proportion of *Ae. tauschii* segments produced more grain under drought stress in the Colby 2018 trial, raising the possibility that the *Ae. tauschii* alleles of *Ppd-D1* also have the potential to protect wheat from the physiological effects of stress that lead to low yield.

Our study reveals that some haplotypes associated with productivity trait variation in the introgression population also exhibit significant pleiotropic effects. While the direction of effects on various traits was largely consistent with the previously reported trade-offs among component traits (Quintero et al., 2018; Griffiths et al., 2015; Reynolds et al., 2017), the combined effects of some introgressed haplotypes were associated with the positive trends in yield. For example, a haplotype contributed by *Ae. tauschii* ssp. *tauschii* at the chromosome 2D haplotype block at 65,964,778 – 66,124,103 bp was associated with increase in grain length, size and number with moderate positive effect on grain yield. At the haplotype block on chromosome 7D located between 14,722,457 – 14,817,138 bp, the Hap0_HW&AeT haplotype shared between hexaploid wheat and *Ae. tauschii* parents and associated with moderate increase in both spikelet number per spike and grain length was also associated with the most significant increase in grain yield. The analyses of pleotropic effects of introgressed haplotypes suggest that these haplotypes on chromosomes 2D and 7D could be utilized in breeding programs to improve yield component traits without negative effects on other productivity traits.

## Conclusions

Imputation of markers from whole genome sequenced reference panels into skim-sequenced inference populations is increasingly becoming a common practice in plant breeding program due to its cost-effectiveness (Happ et al., 2019; Jessen et al., 2020). Our study demonstrates the utility of this strategy for detecting introgression in the wheat genome and contributes to developing genomic resources for deploying wild relative diversity in wheat breeding programs. We show that the haplotype-based analysis of trait variation in this population has the potential to improve our knowledge on the genetic effects of introgressed diversity on productivity traits and identify novel haplotypes for improving yield potential in wheat.

## Supporting information

Summary of supplemental files

## 5 Conflict of Interest

The authors declare that the research was conducted in the absence of any commercial or financial relationships that could be construed as a potential conflict of interest.

## 6 Author Contributions

AKF and MC generated the *Ae. tauschii* introgression population. MN, EA, MC, RA, BB, WW, DD, ZY, YG, FH, KWJ and AKF phenotyped the introgression population. MN, BB and AA generated the genotyping data. AA performed next-generation sequencing of parental lines and introgression population; MN and FH performed the bioinformatical analyses of data. MN performed the statistical analysis; EA conceived the idea and interpreted results; MN and EA wrote the manuscript. All co-authors contributed to the manuscript revision, read and approved the submitted version.

## 7 Funding

This research was supported by the Agriculture and Food Research Initiative Competitive Grants 2017–67007-25939 (WheatCAP) and 2020-68013-30905 from the USDA National Institute of Food and Agriculture, and by the International Wheat Yield Partnership (IWYP).

## 8 Acknowledgments

We would like to thank Mary Guttieri for providing access to the USDA ARS seed processing facility.

## Notes

### Competing Interest Statement

The authors have declared no competing interest.

### Summary of Updates

The current revision includes a detailed analysis of SNP diversity at the Ppd-D1 locus including their effect on amino acid sequences using only whole-genome sequencing data generated for all wheat and Ae. tauschii parental lines. This analysis revealed eight (8) haplotypes of the Ppd-D1 gene, with one haplotype shared by all hexaploid wheat parents, and remaining seven haplotypes found in Ae. tauschii. These haplotypes and variation of surrounding regions are shown in the revised Figure 5A. We used SNPs flanking the Ppd-D1 locus but also present in the introgression population to infer the parental Ppd-D1 haplotypes in the introgression population and assessed their effects on spikelet number and heading date. The results of the Ppd-D1 haplotype effects are provided in Table 4. Finally, we demonstrate that SNPs and haplotypes significantly associated with variation in spikelet number and heading date are also associated with the haplotypes of the Ppd-D1 locus. We show that the Ppd-D1 haplotypes from Ae. tauschii are linked with SNPs and haplotypes from Ae. tauschii that are significantly associated with spikelet number and heading date. The results of these analyses as well as phenotypic effects of GWAS SNPs and Ppd-D1 haplotypes are presented in the updated Figure 5B.

