## Supplemental Figure 1 for "The haplotype-based analysis of *Aegilops tauschii* introgression into hard red winter wheat and its impact on productivity traits"

### Slide 1
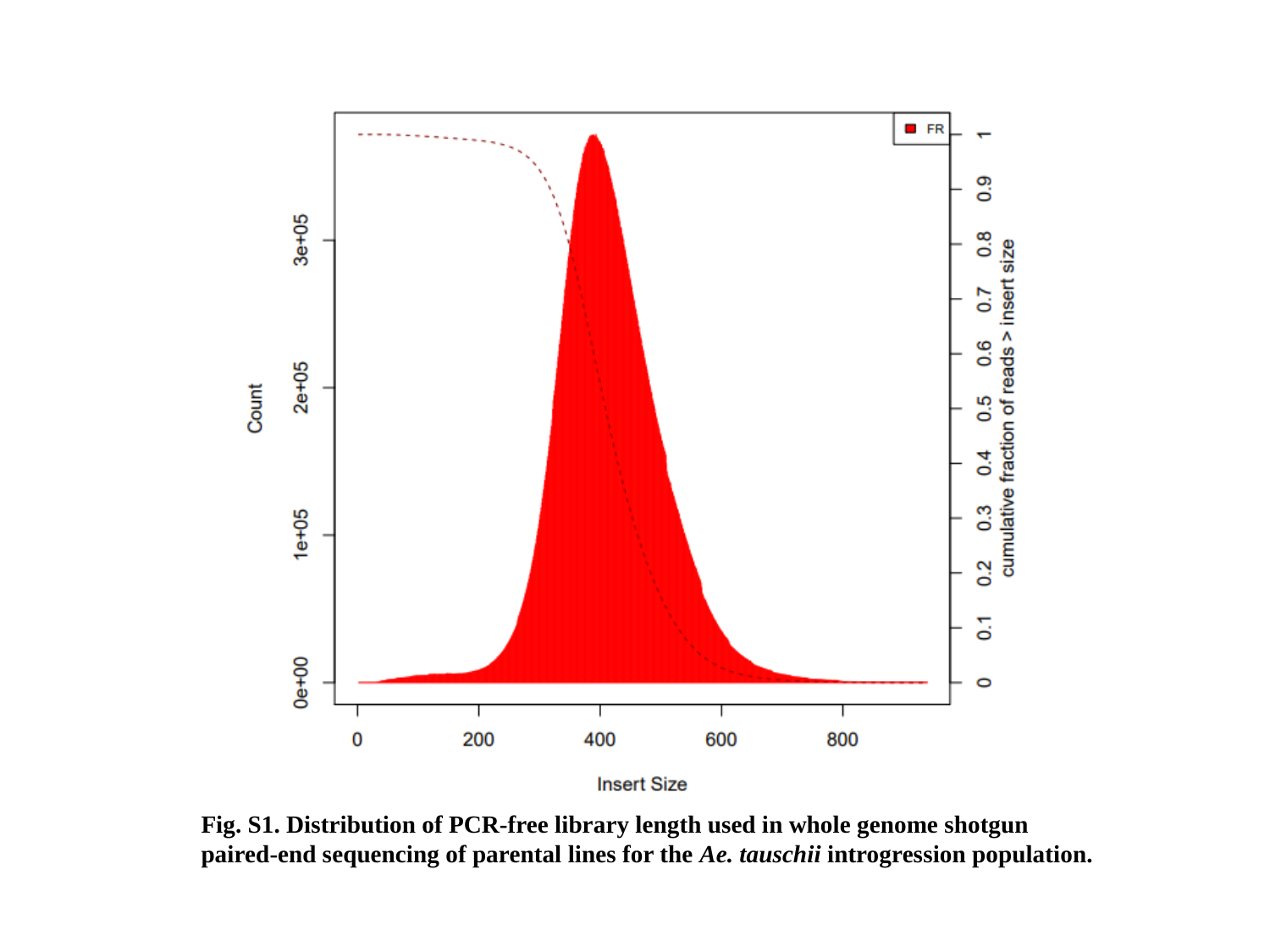

Fig. S1. Distribution of PCR-free library length used in whole genome shotgun paired-end sequencing of parental lines for the Ae. tauschii introgression population.
