## Supplemental Figure 2 for "The haplotype-based analysis of *Aegilops tauschii* introgression into hard red winter wheat and its impact on productivity traits"

### Slide 1
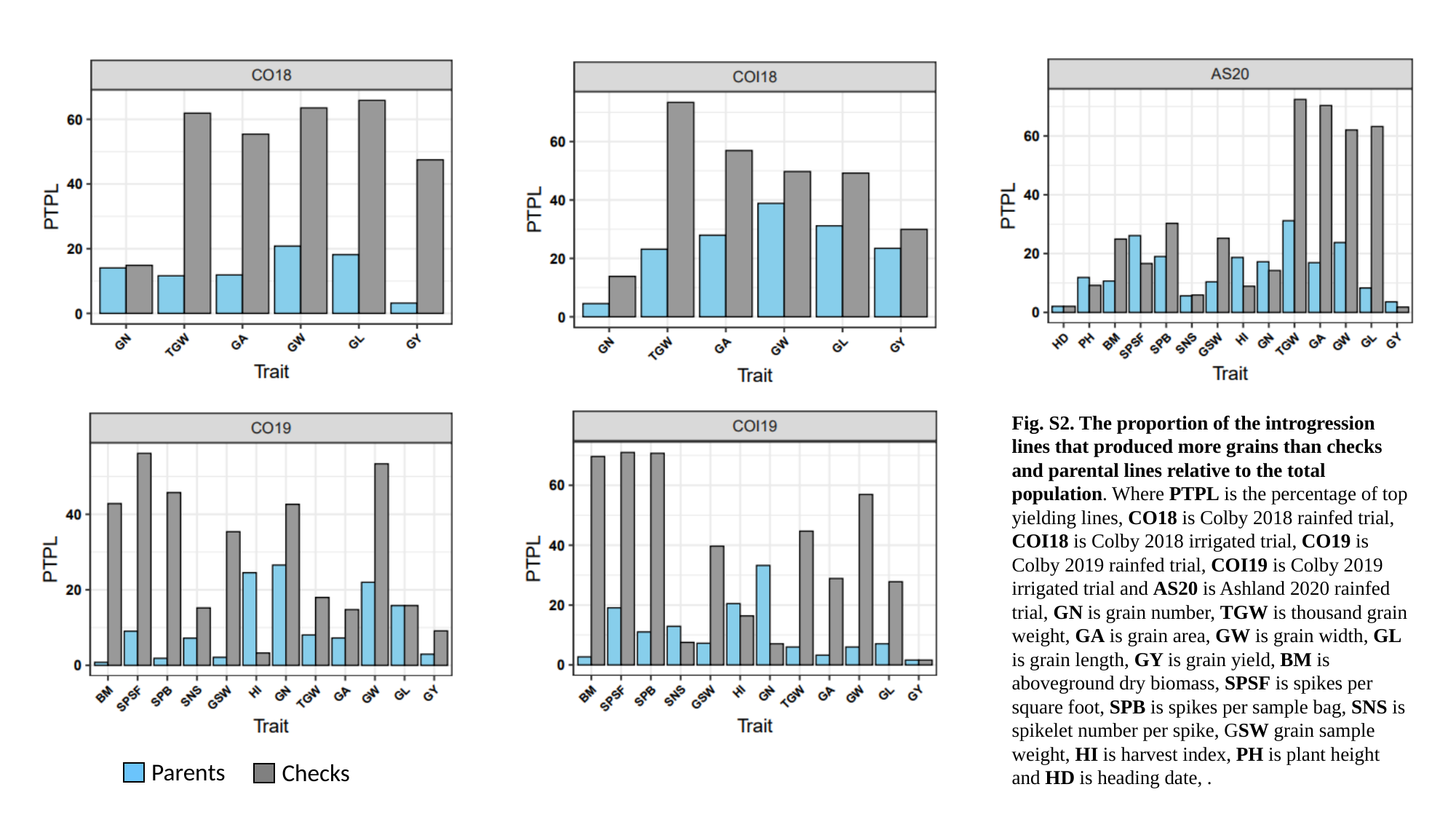

Fig. S2. The proportion of the introgression lines that produced more grains than checks and parental lines relative to the total population. Where PTPL is the percentage of top yielding lines, CO18 is Colby 2018 rainfed trial, COI18 is Colby 2018 irrigated trial, CO19 is Colby 2019 rainfed trial, COI19 is Colby 2019 irrigated trial and AS20 is Ashland 2020 rainfed trial, GN is grain number, TGW is thousand grain weight, GA is grain area, GW is grain width, GL is grain length, GY is grain yield, BM is aboveground dry biomass, SPSF is spikes per square foot, SPB is spikes per sample bag, SNS is spikelet number per spike, GSW grain sample weight, HI is harvest index, PH is plant height and HD is heading date, .
Parents
Checks
