## Supplemental File 4 for "The haplotype-based analysis of *Aegilops tauschii* introgression into hard red winter wheat and its impact on productivity traits"

### Slide 1
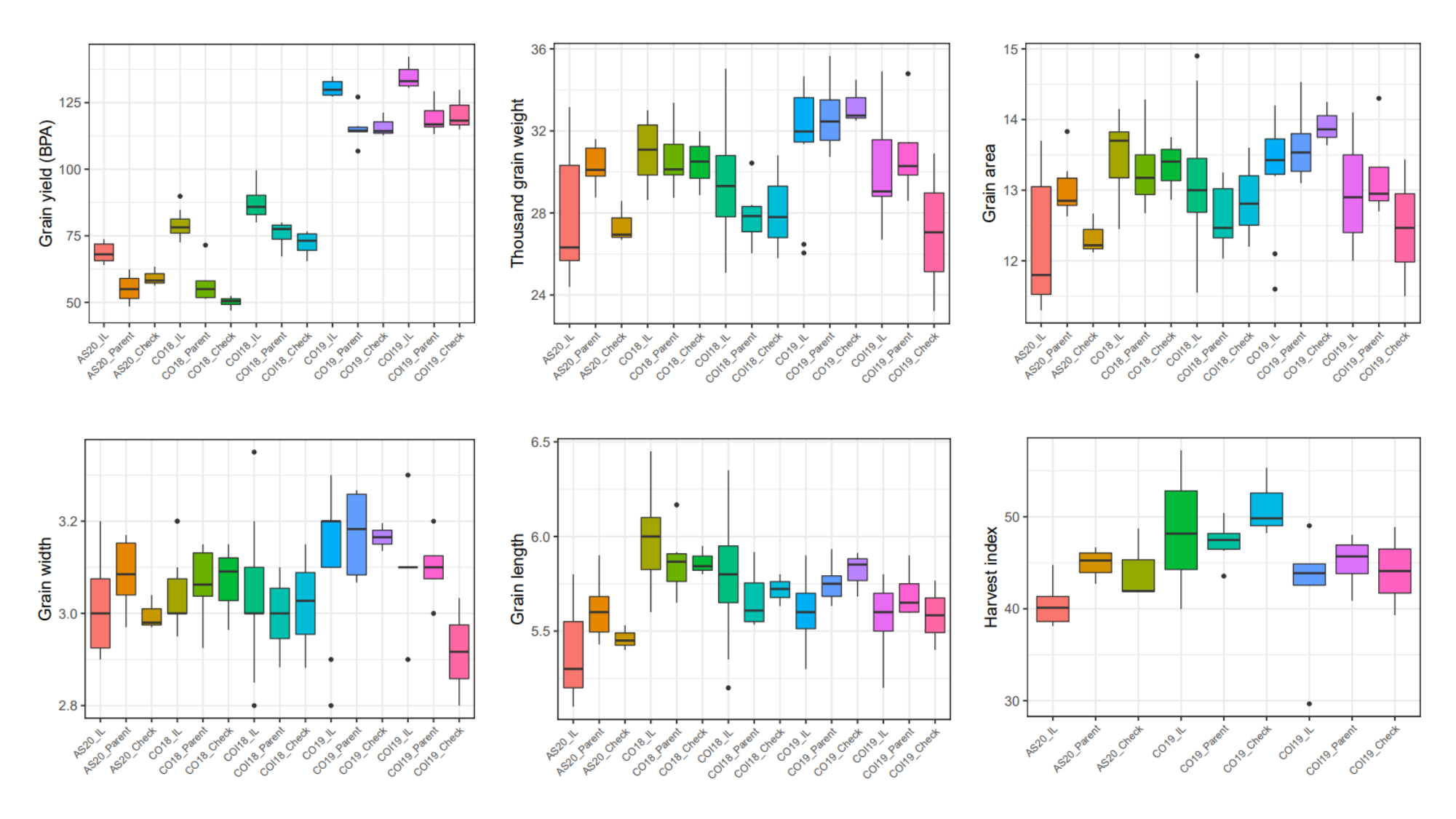

### Slide 2
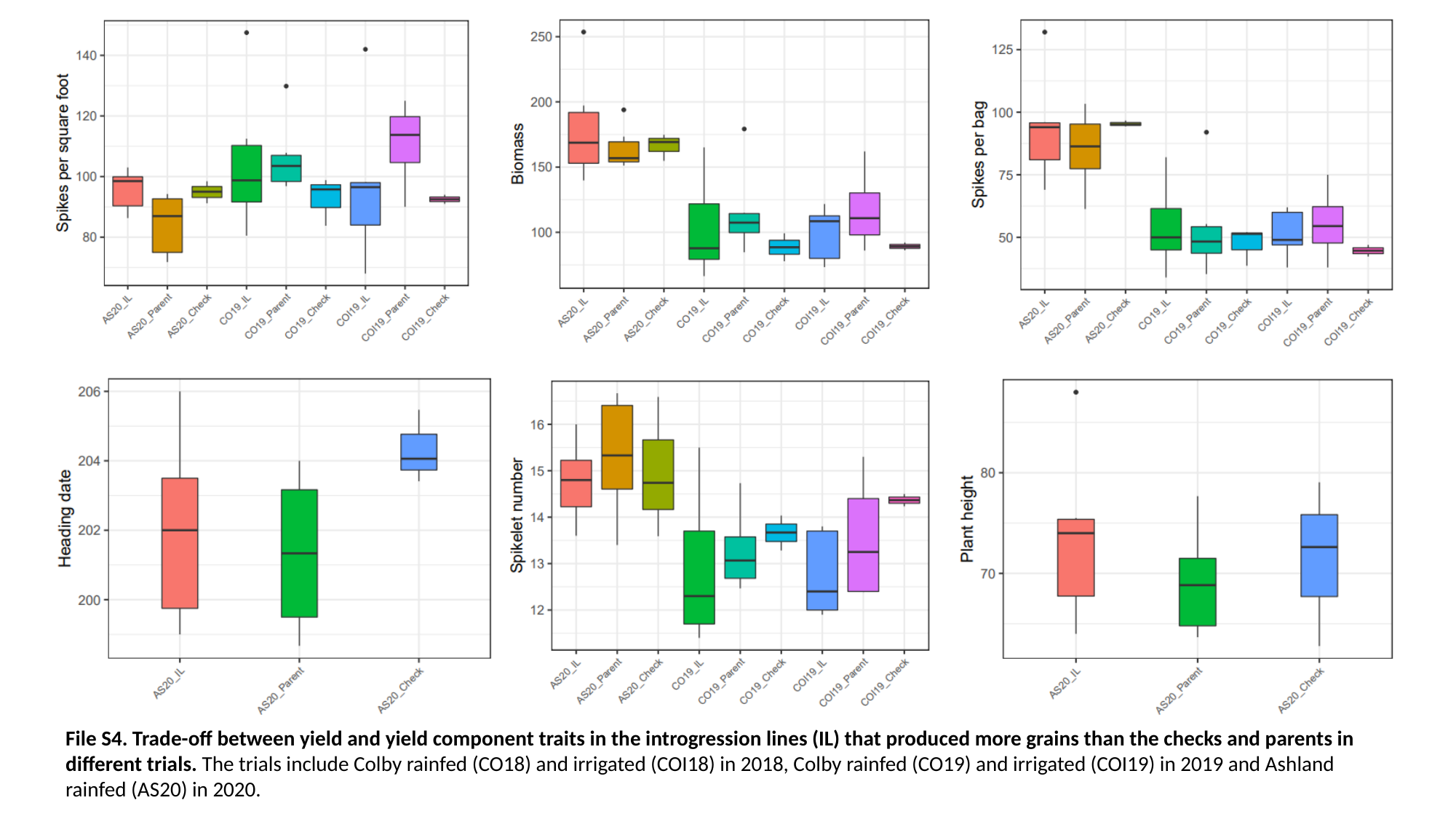

File S4. Trade-off between yield and yield component traits in the introgression lines (IL) that produced more grains than the checks and parents in different trials. The trials include Colby rainfed (CO18) and irrigated (COI18) in 2018, Colby rainfed (CO19) and irrigated (COI19) in 2019 and Ashland rainfed (AS20) in 2020.
