## Supplemental File 5 for "The haplotype-based analysis of *Aegilops tauschii* introgression into hard red winter wheat and its impact on productivity traits"

### Slide 1
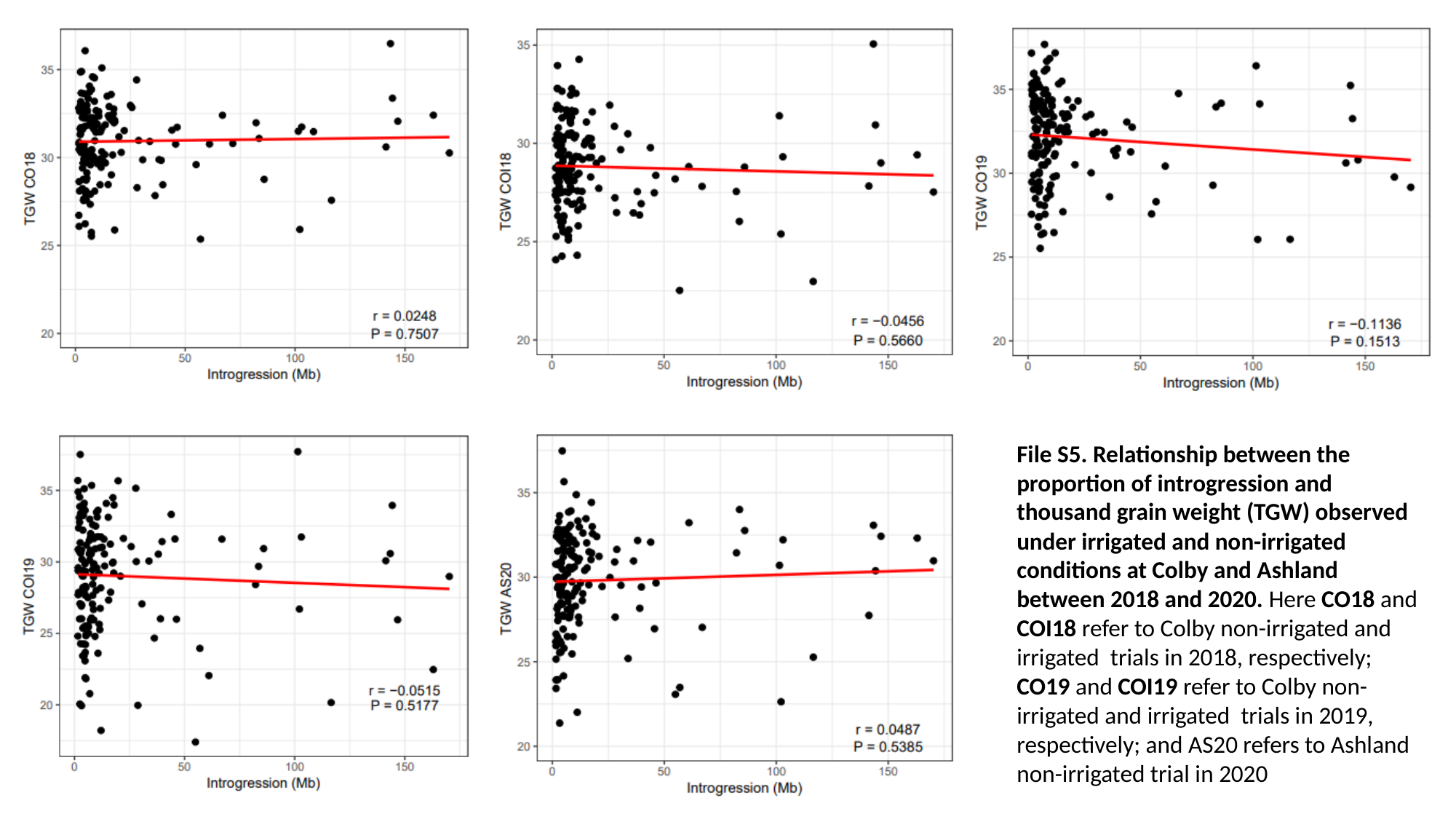

File S5. Relationship between the proportion of introgression and thousand grain weight (TGW) observed under irrigated and non-irrigated conditions at Colby and Ashland between 2018 and 2020. Here CO18 and COI18 refer to Colby non-irrigated and irrigated trials in 2018, respectively; CO19 and COI19 refer to Colby non-irrigated and irrigated trials in 2019, respectively; and AS20 refers to Ashland non-irrigated trial in 2020

### Slide 2
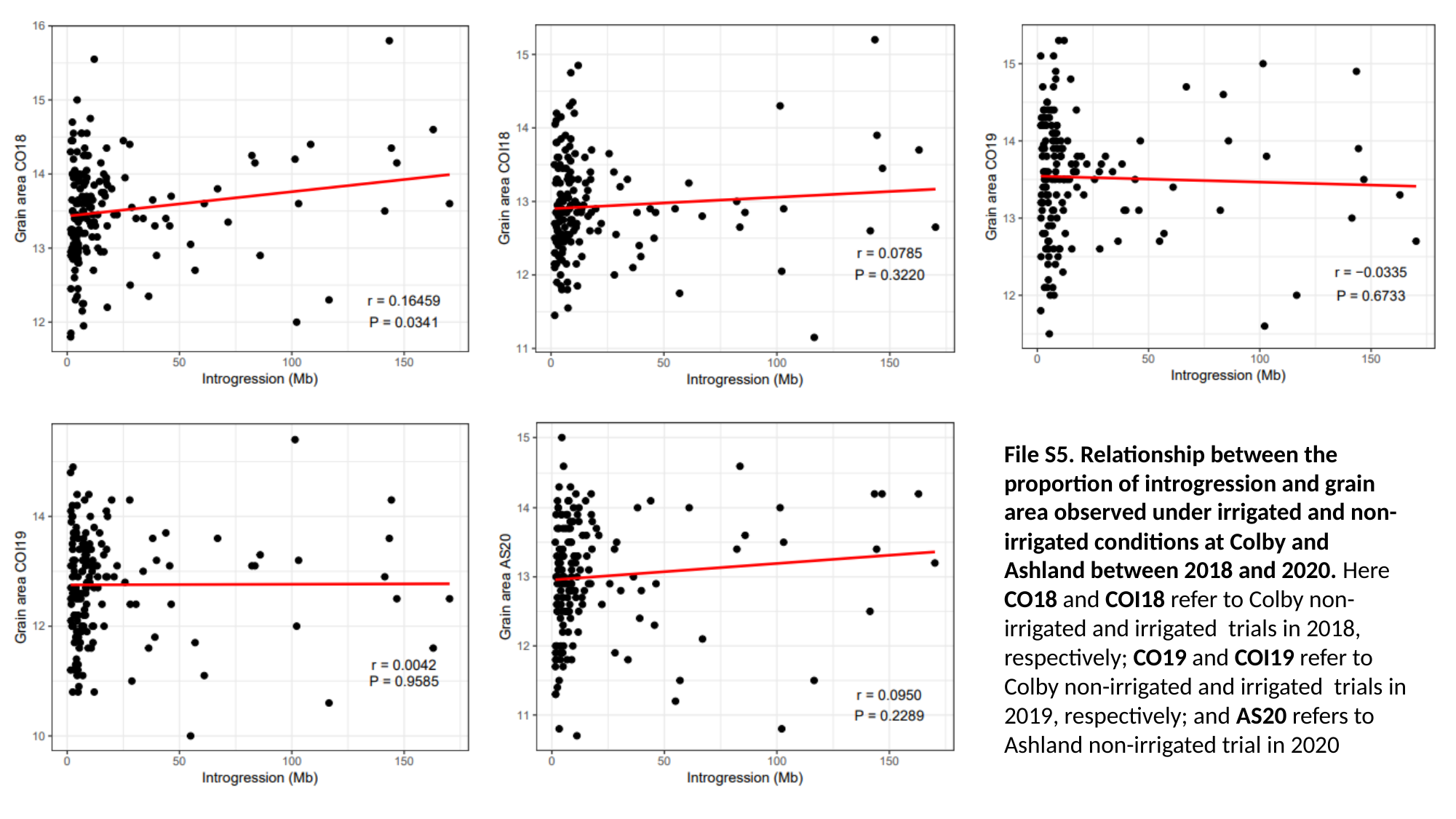

File S5. Relationship between the proportion of introgression and grain area observed under irrigated and non-irrigated conditions at Colby and Ashland between 2018 and 2020. Here CO18 and COI18 refer to Colby non-irrigated and irrigated trials in 2018, respectively; CO19 and COI19 refer to Colby non-irrigated and irrigated trials in 2019, respectively; and AS20 refers to Ashland non-irrigated trial in 2020

### Slide 3
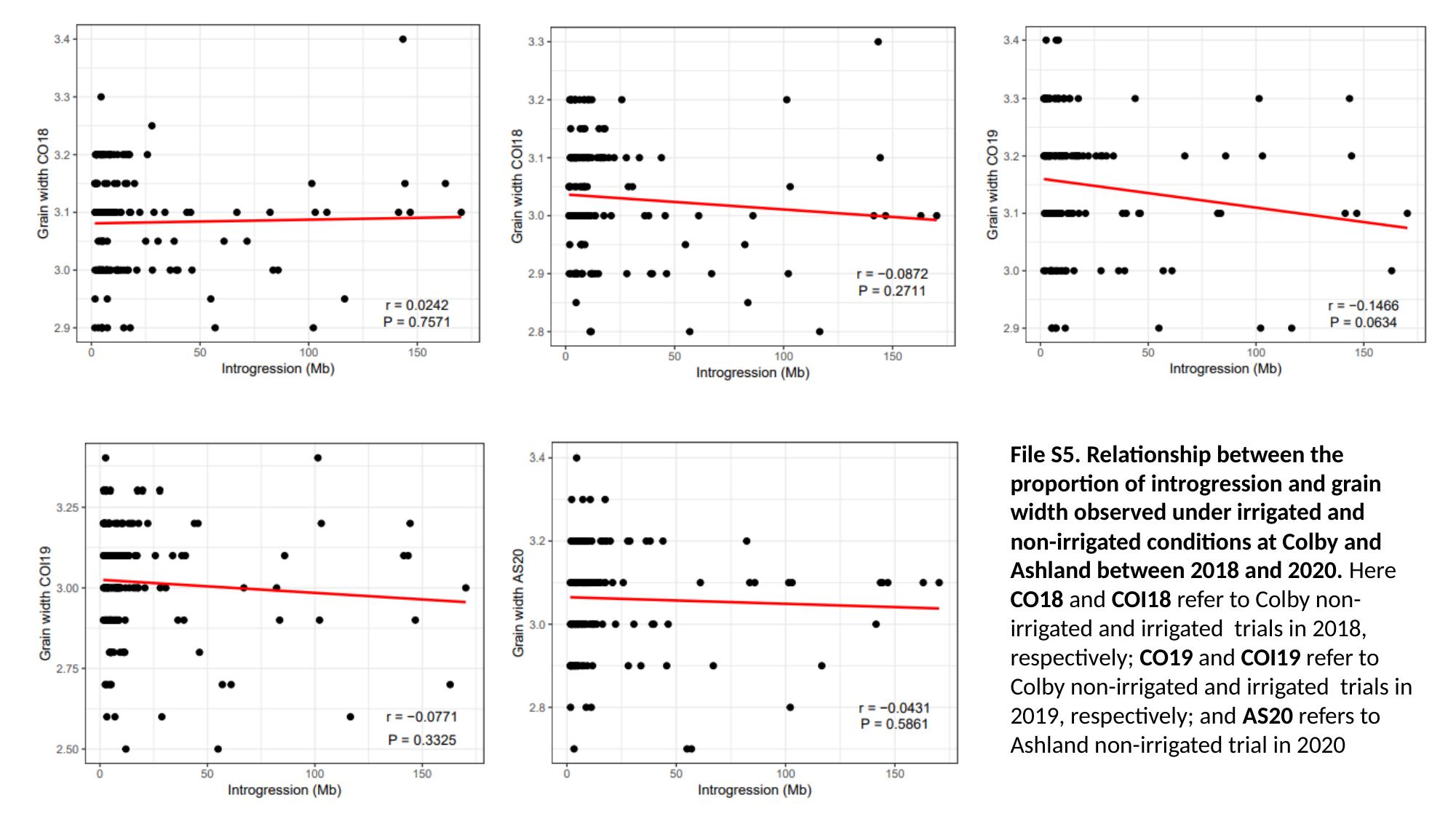

File S5. Relationship between the proportion of introgression and grain width observed under irrigated and non-irrigated conditions at Colby and Ashland between 2018 and 2020. Here CO18 and COI18 refer to Colby non-irrigated and irrigated trials in 2018, respectively; CO19 and COI19 refer to Colby non-irrigated and irrigated trials in 2019, respectively; and AS20 refers to Ashland non-irrigated trial in 2020

### Slide 4
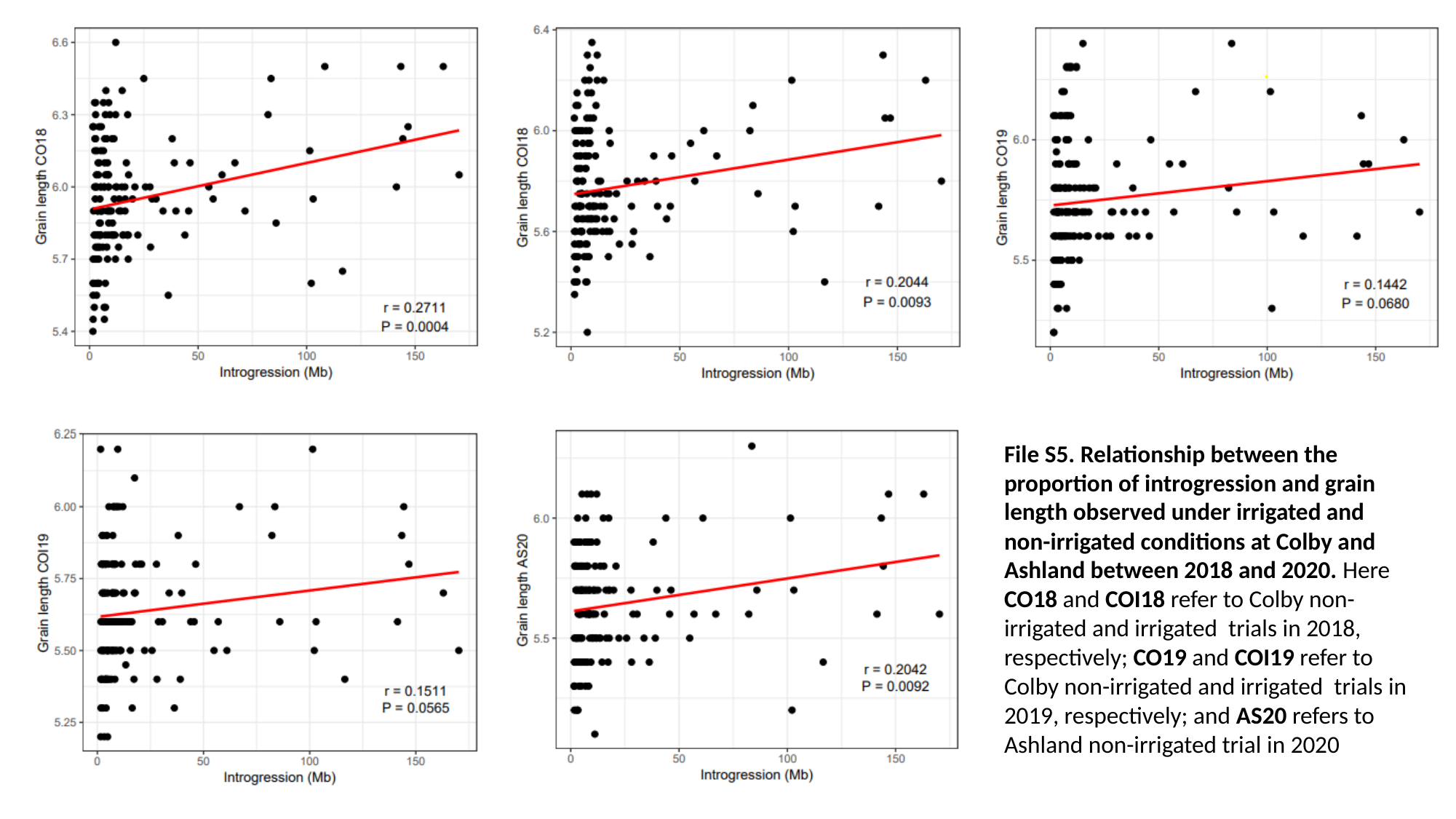

File S5. Relationship between the proportion of introgression and grain length observed under irrigated and non-irrigated conditions at Colby and Ashland between 2018 and 2020. Here CO18 and COI18 refer to Colby non-irrigated and irrigated trials in 2018, respectively; CO19 and COI19 refer to Colby non-irrigated and irrigated trials in 2019, respectively; and AS20 refers to Ashland non-irrigated trial in 2020

### Slide 5
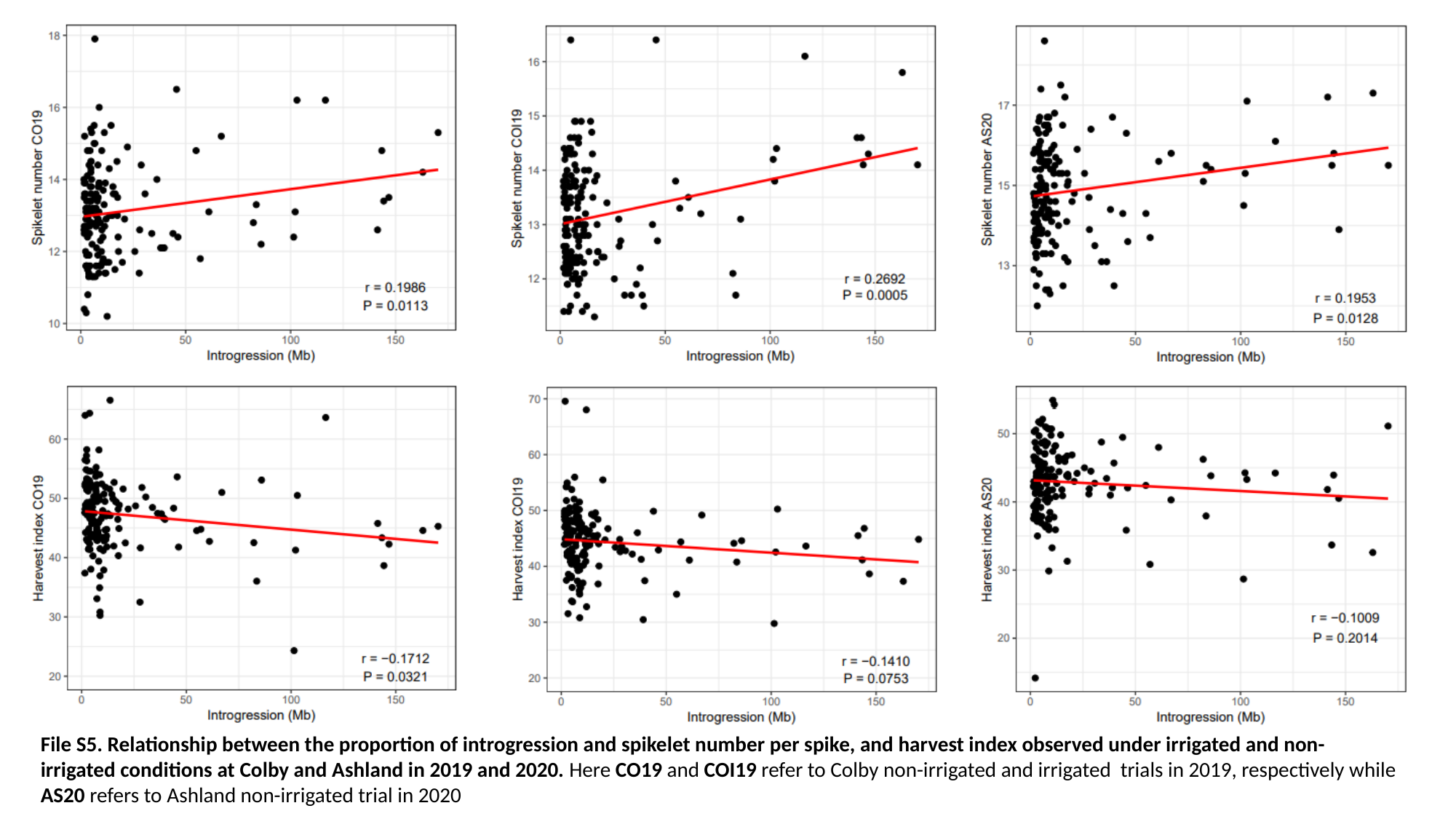

File S5. Relationship between the proportion of introgression and spikelet number per spike, and harvest index observed under irrigated and non-irrigated conditions at Colby and Ashland in 2019 and 2020. Here CO19 and COI19 refer to Colby non-irrigated and irrigated trials in 2019, respectively while AS20 refers to Ashland non-irrigated trial in 2020

### Slide 6
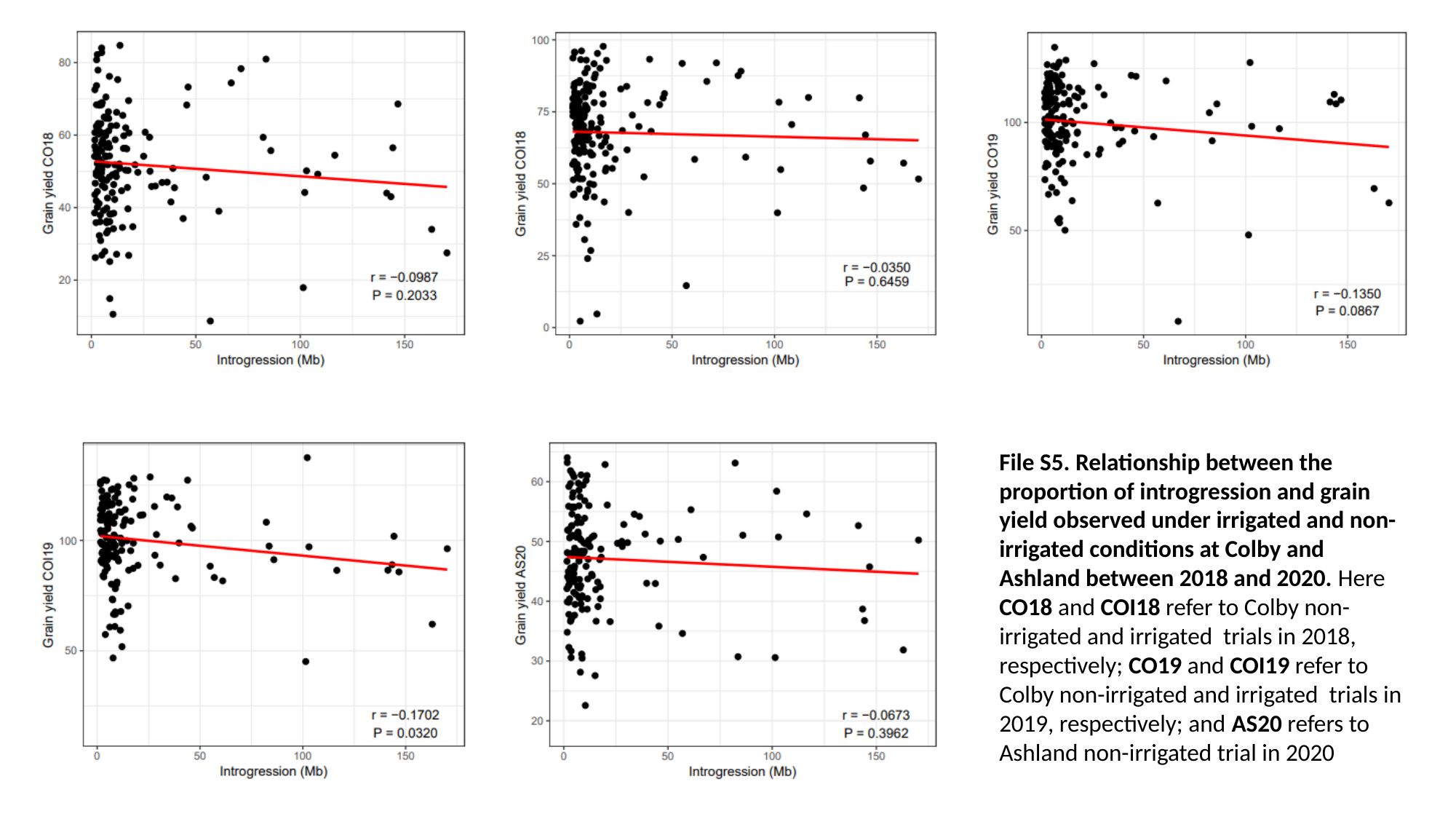

File S5. Relationship between the proportion of introgression and grain yield observed under irrigated and non-irrigated conditions at Colby and Ashland between 2018 and 2020. Here CO18 and COI18 refer to Colby non-irrigated and irrigated trials in 2018, respectively; CO19 and COI19 refer to Colby non-irrigated and irrigated trials in 2019, respectively; and AS20 refers to Ashland non-irrigated trial in 2020
