## Supplemental File 7 for "The haplotype-based analysis of *Aegilops tauschii* introgression into hard red winter wheat and its impact on productivity traits"

### Slide 1
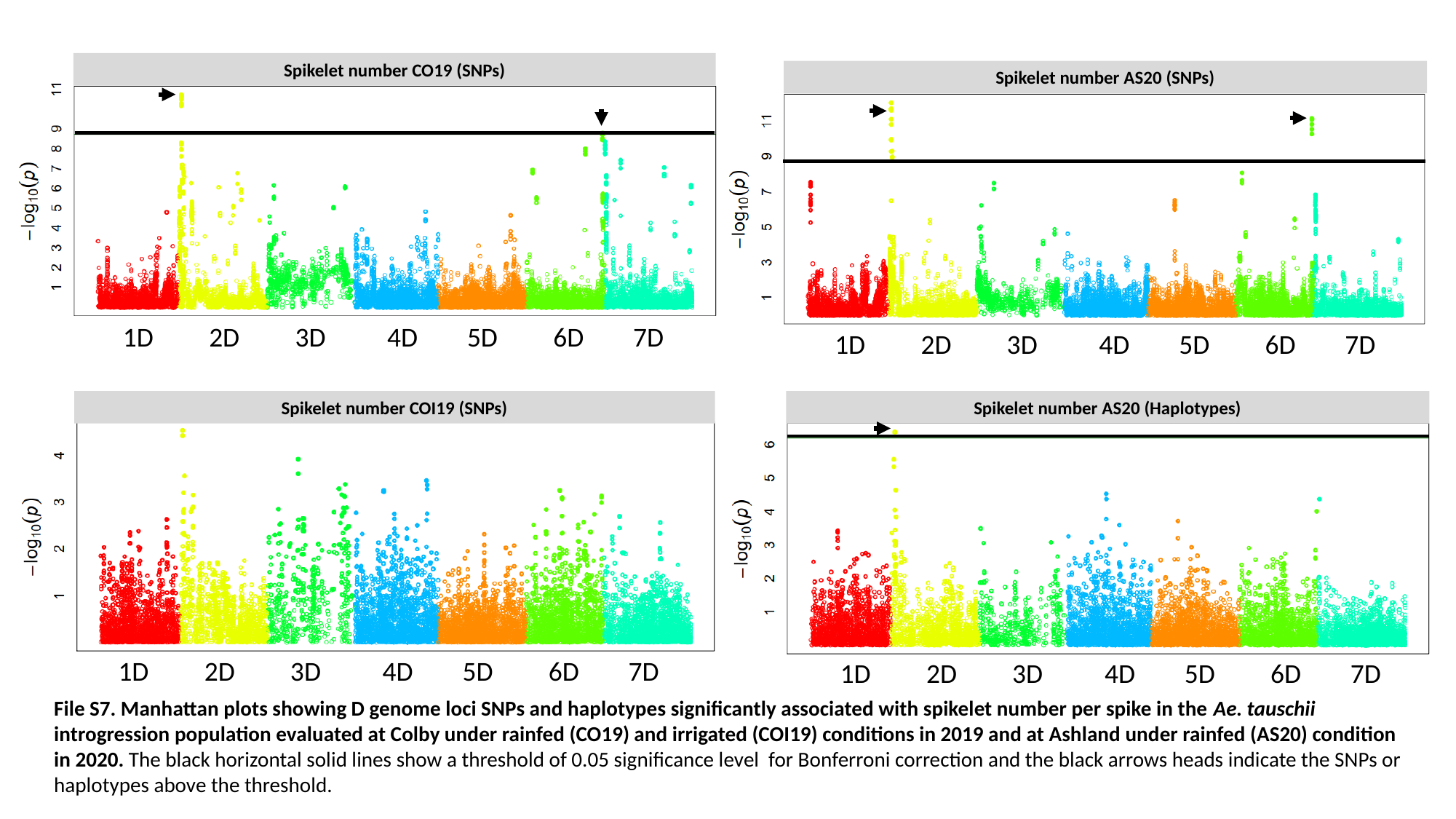

Spikelet number CO19 (SNPs)
 1D 2D 3D 4D 5D 6D 7D
Spikelet number COI19 (SNPs)
 1D 2D 3D 4D 5D 6D 7D
Spikelet number AS20 (SNPs)
 1D 2D 3D 4D 5D 6D 7D
Spikelet number AS20 (Haplotypes)
 1D 2D 3D 4D 5D 6D 7D
File S7. Manhattan plots showing D genome loci SNPs and haplotypes significantly associated with spikelet number per spike in the Ae. tauschii introgression population evaluated at Colby under rainfed (CO19) and irrigated (COI19) conditions in 2019 and at Ashland under rainfed (AS20) condition in 2020. The black horizontal solid lines show a threshold of 0.05 significance level for Bonferroni correction and the black arrows heads indicate the SNPs or haplotypes above the threshold.

### Slide 2
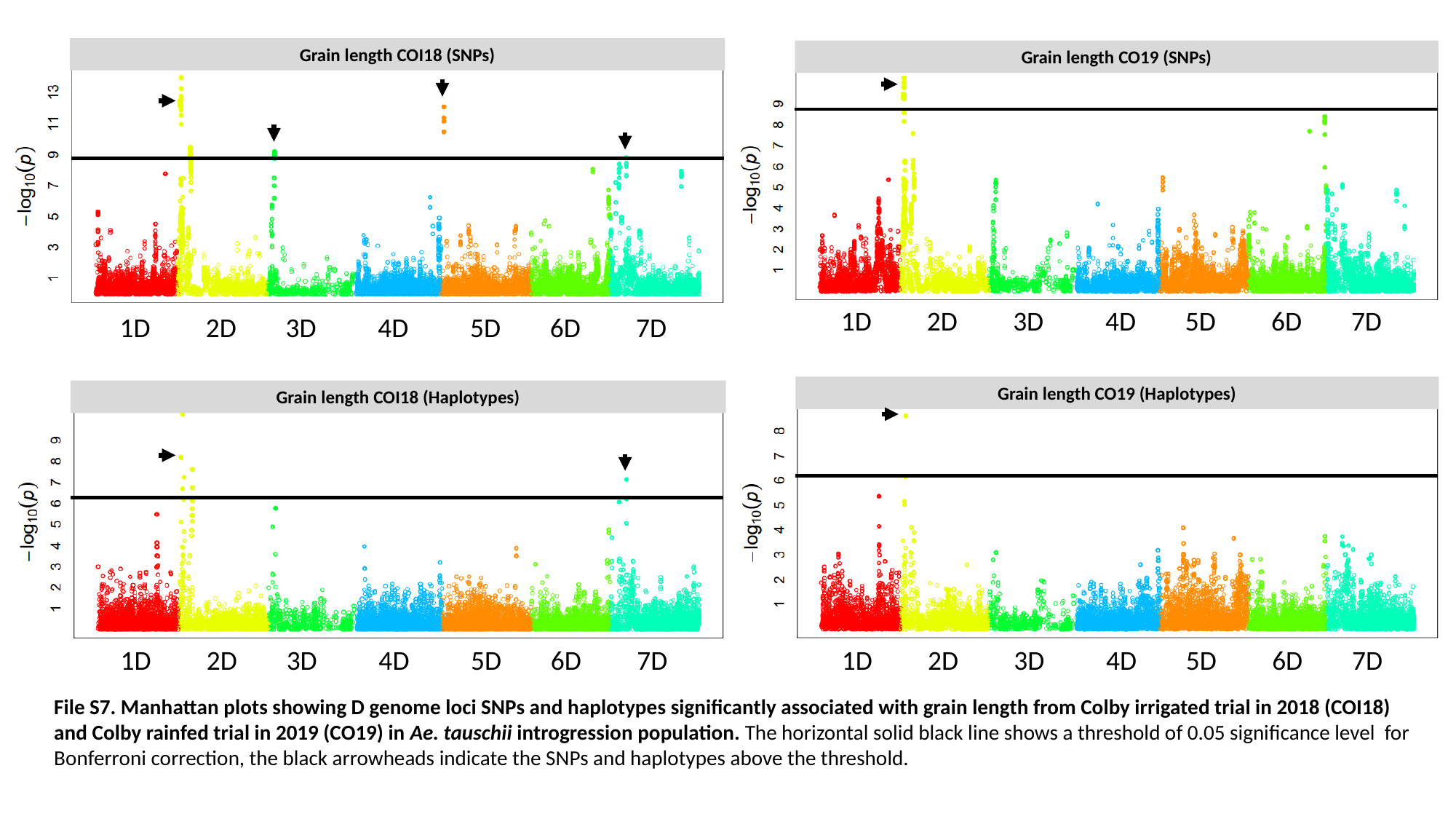

Grain length COI18 (SNPs)
 1D 2D 3D 4D 5D 6D 7D
Grain length CO19 (SNPs)
 1D 2D 3D 4D 5D 6D 7D
Grain length CO19 (Haplotypes)
 1D 2D 3D 4D 5D 6D 7D
Grain length COI18 (Haplotypes)
 1D 2D 3D 4D 5D 6D 7D
File S7. Manhattan plots showing D genome loci SNPs and haplotypes significantly associated with grain length from Colby irrigated trial in 2018 (COI18) and Colby rainfed trial in 2019 (CO19) in Ae. tauschii introgression population. The horizontal solid black line shows a threshold of 0.05 significance level for Bonferroni correction, the black arrowheads indicate the SNPs and haplotypes above the threshold.

### Slide 3
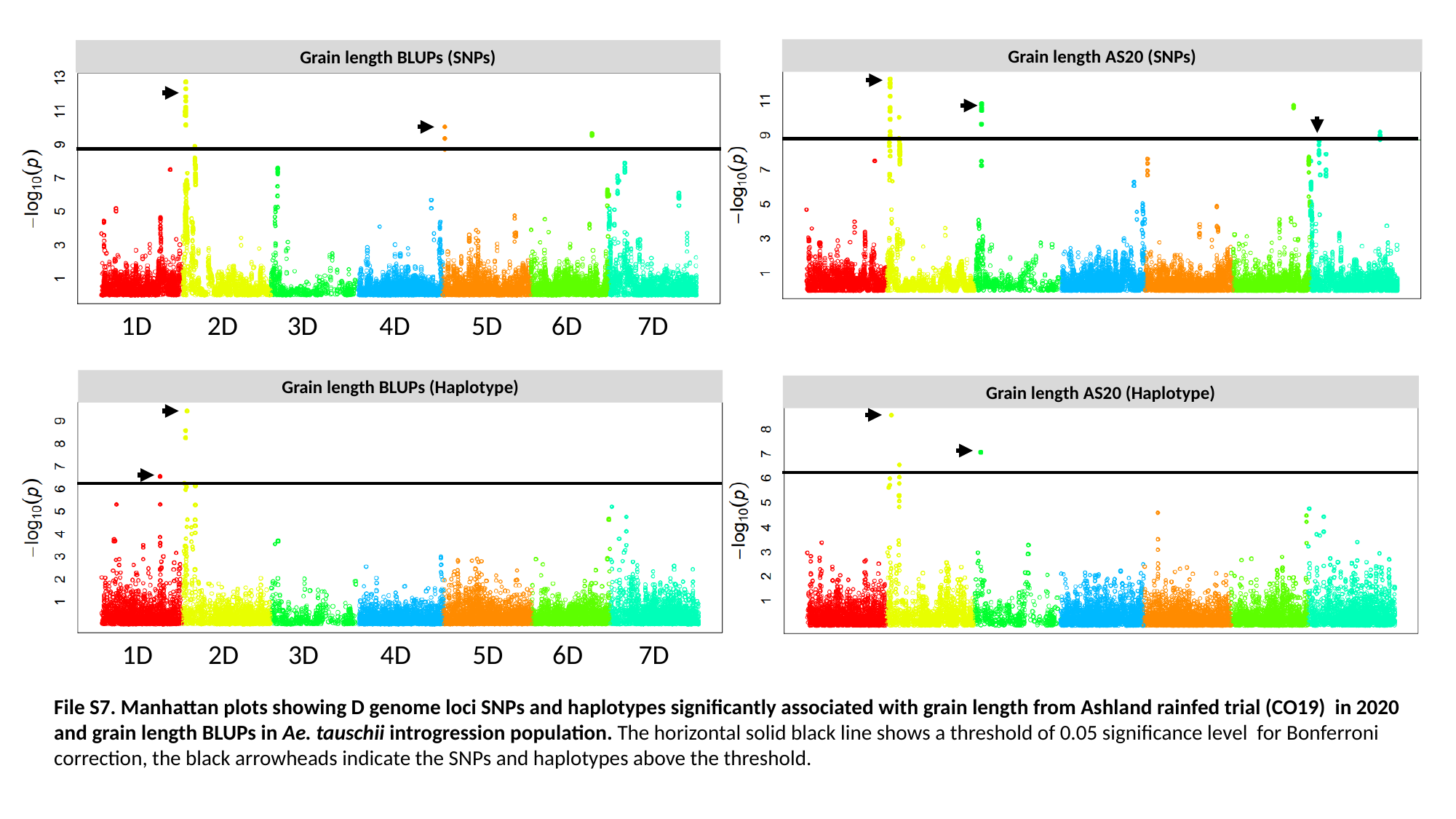

Grain length AS20 (SNPs)
Grain length AS20 (Haplotype)
Grain length BLUPs (SNPs)
 1D 2D 3D 4D 5D 6D 7D
Grain length BLUPs (Haplotype)
 1D 2D 3D 4D 5D 6D 7D
File S7. Manhattan plots showing D genome loci SNPs and haplotypes significantly associated with grain length from Ashland rainfed trial (CO19) in 2020 and grain length BLUPs in Ae. tauschii introgression population. The horizontal solid black line shows a threshold of 0.05 significance level for Bonferroni correction, the black arrowheads indicate the SNPs and haplotypes above the threshold.

### Slide 4
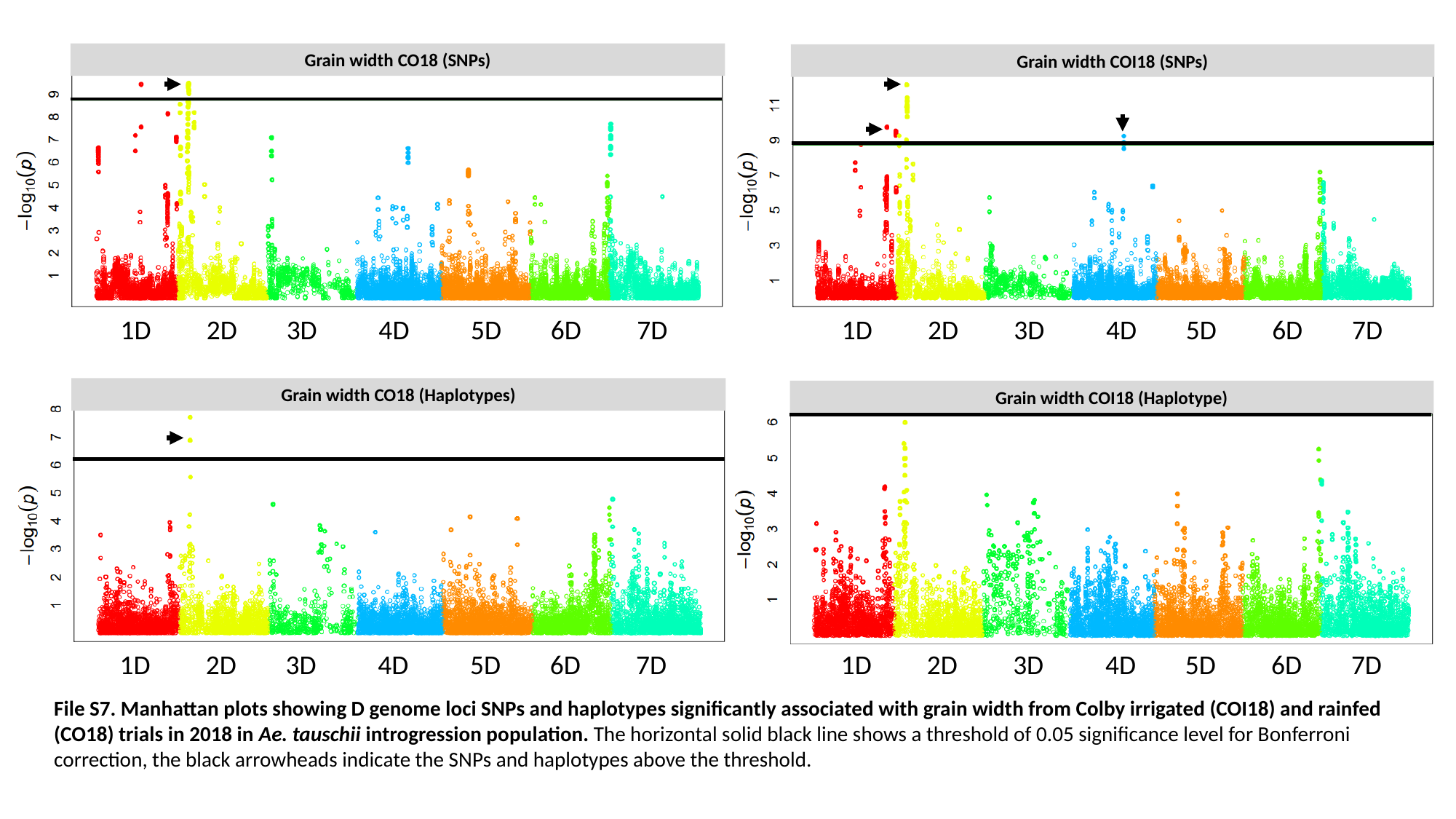

Grain width CO18 (SNPs)
 1D 2D 3D 4D 5D 6D 7D
Grain width COI18 (SNPs)
 1D 2D 3D 4D 5D 6D 7D
Grain width CO18 (Haplotypes)
 1D 2D 3D 4D 5D 6D 7D
Grain width COI18 (Haplotype)
 1D 2D 3D 4D 5D 6D 7D
File S7. Manhattan plots showing D genome loci SNPs and haplotypes significantly associated with grain width from Colby irrigated (COI18) and rainfed (CO18) trials in 2018 in Ae. tauschii introgression population. The horizontal solid black line shows a threshold of 0.05 significance level for Bonferroni correction, the black arrowheads indicate the SNPs and haplotypes above the threshold.

### Slide 5
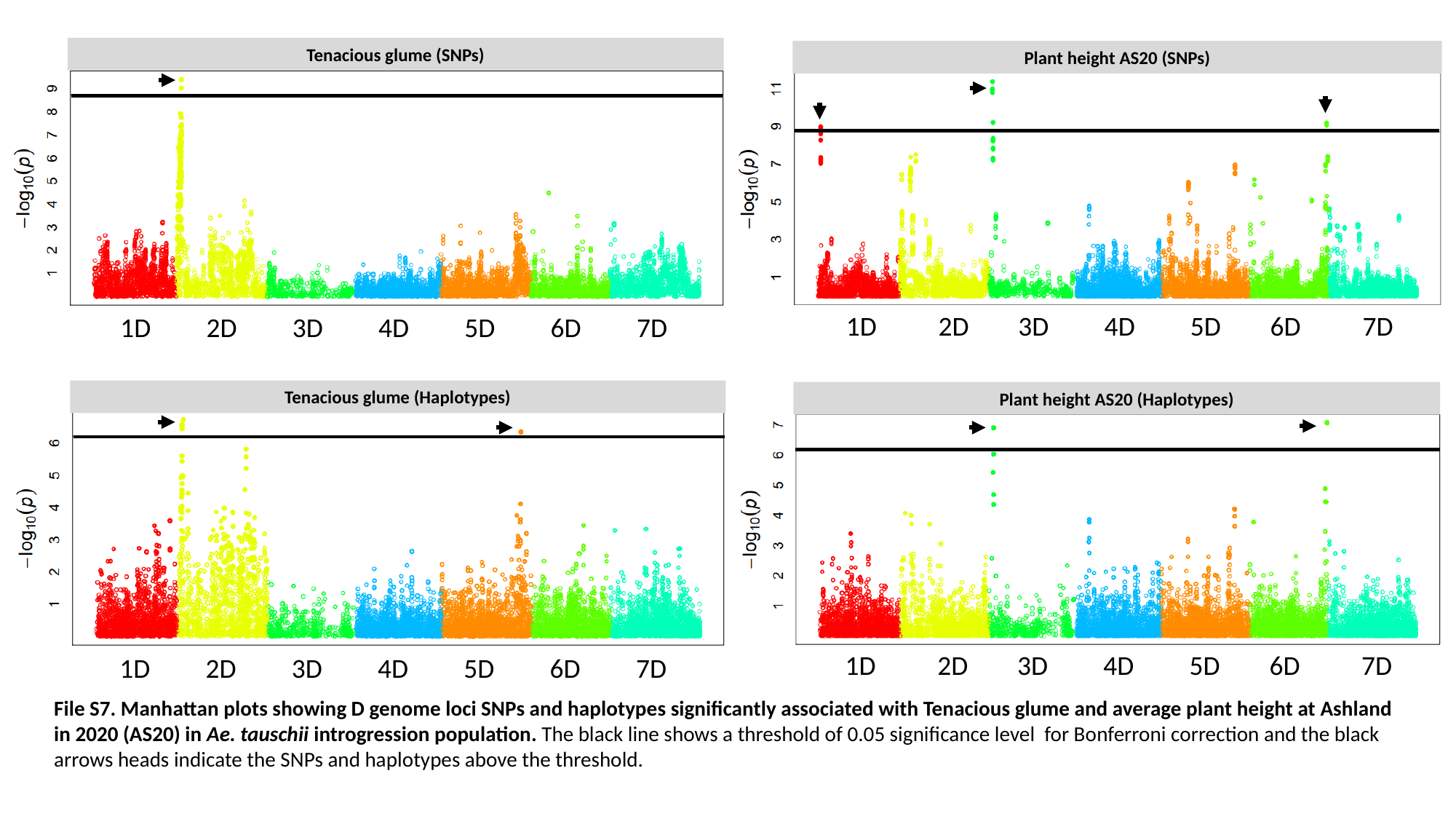

Tenacious glume (SNPs)
 1D 2D 3D 4D 5D 6D 7D
Plant height AS20 (SNPs)
 1D 2D 3D 4D 5D 6D 7D
Tenacious glume (Haplotypes)
 1D 2D 3D 4D 5D 6D 7D
Plant height AS20 (Haplotypes)
 1D 2D 3D 4D 5D 6D 7D
File S7. Manhattan plots showing D genome loci SNPs and haplotypes significantly associated with Tenacious glume and average plant height at Ashland in 2020 (AS20) in Ae. tauschii introgression population. The black line shows a threshold of 0.05 significance level for Bonferroni correction and the black arrows heads indicate the SNPs and haplotypes above the threshold.

### Slide 6
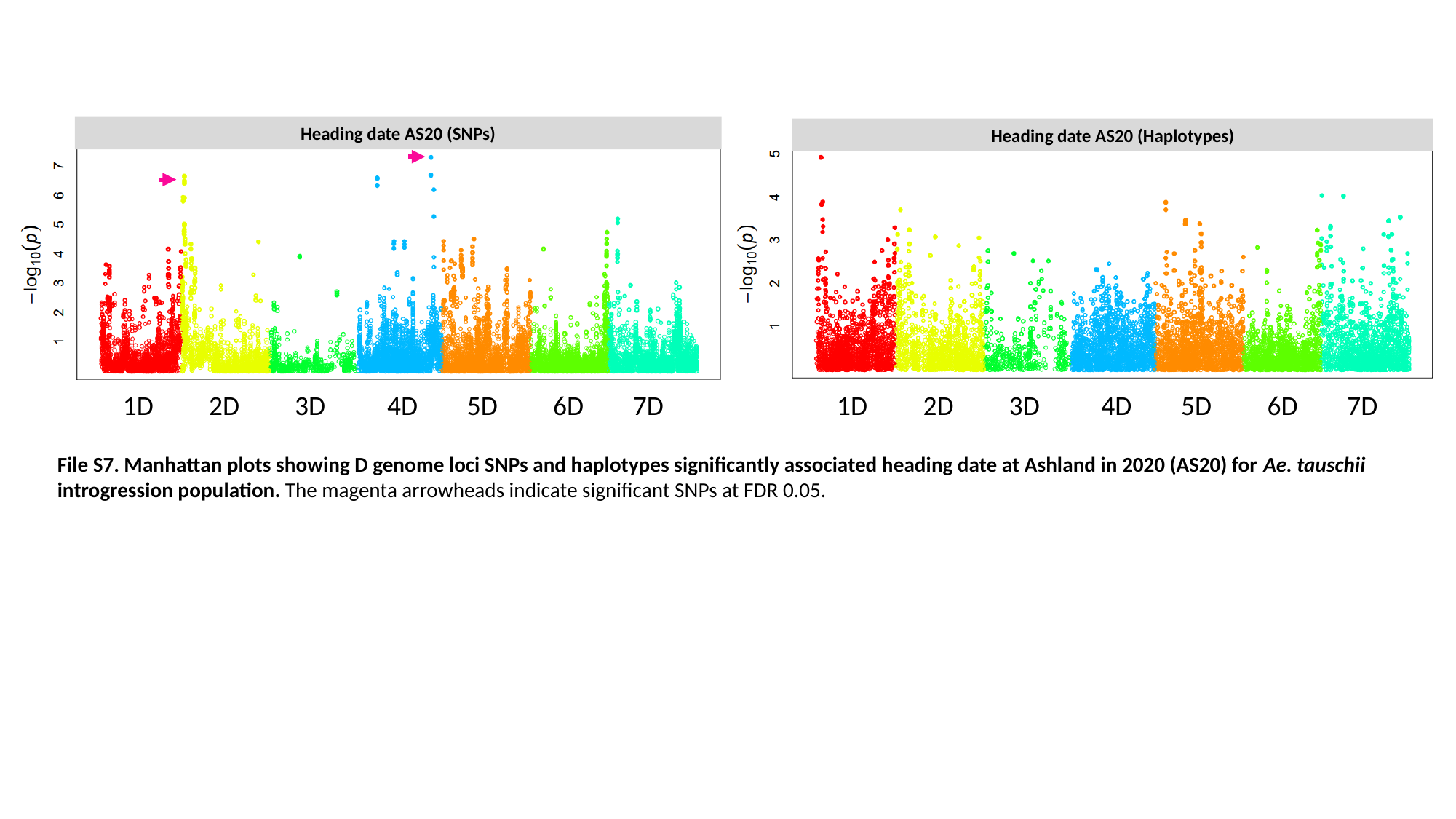

Heading date AS20 (SNPs)
 1D 2D 3D 4D 5D 6D 7D
Heading date AS20 (Haplotypes)
 1D 2D 3D 4D 5D 6D 7D
File S7. Manhattan plots showing D genome loci SNPs and haplotypes significantly associated heading date at Ashland in 2020 (AS20) for Ae. tauschii introgression population. The magenta arrowheads indicate significant SNPs at FDR 0.05.
