## Supplementary material for "The haplotype-based analysis of *Aegilops tauschii* introgression into hard red winter wheat and its impact on productivity traits": Summary of supplemental files

### 1 Supplementary Files

**File S1.** Estimation of yield stability across treatments (rainfed and irrigated), years (2018 - 2020) and locations (Ashaland and Colby) using parametric and non-parametric methods.

**File S2.** Estimation of harvest index stability across treatments (rainfed and irrigated), years (2018 - 2020) and locations (Ashaland and Colby) using parametric and non-parametric methods.

**File S3.** Pearson's correlation coefficients between traits within treatments per year and location in the *Ae. tauschii* introgression population.

**File S4.** Trade-off between yield and yield component traits in the introgression lines (IL) that produced more grains than the checks and parents in different trials. The trials include Colby rainfed (CO18) and irrigated (COI18) in 2018, Colby rainfed (CO19) and irrigated (COI19) in 2019 and Ashland rainfed (AS20) in 2020.

**File S5.** Relationship between the proportion of introgression and the phenotypes in the *Ae. tauschii* introgression population.

**File S6.** *Ae. tauschii* haplotypes in the introgression lines derived from KanMark (hexaploid wheat) x TA1642 (*Ae. tauschii* ssp. *strangulata*) (FAM93) and Danby (hexaploid wheat) x TA1642 (*Ae. tauschii* ssp. *tauschii*) (FAM97).

**File S7.** Manhattan plots showing the D genome loci with SNPs and haplotypes that are significantly associated with different traits evaluated in the *Ae. tauschii* introgression population.

**File S8.** Verification of haplotypes associated with spikelet number and heading date in the *Ae. tauschii* introgression population based on the allelic effect of the SNPs observed in the introgression lines with extreme phenotype.

### 2 Supplementary Figures and Tables

#### 2.1 Supplementary Figures

**Figure S1.** Distribution of PCR-free library length used in whole genome shotgun paired-end sequencing of parental lines for the *Ae. tauschii* introgression population.

**Figure S2.** The proportion of the introgression lines that produced more grains than checks and parental lines relative to the total population. Where **PTPL** is the percentage of top yielding lines, **CO18** is Colby 2018 rainfed trial, **COI18** is Colby 2018 irrigated trial, **CO19** is Colby 2019 rainfed trial, **COI19** is Colby 19 irrigated trial and **AS20** is Ashland rainfed trial, **GN** is grain number, **TGW** is thousand grain weight, **BM** is aboveground dry biomass, **SPSF** is spikes per square foot, **SPB** is spikes per sample bag, **SNS** is spikelet number per spike, **GSW** grain sample weight, **HI** is harvest index, **PH** is plant height and **HD** is heading date.

#### 2.2 Supplementary Tables

**Table S1.** Genomic intervals containing significant SNP-trait and haplotype-trait associations for spikelet number per spike (SNS), grain length (GL) and grain width (GW) in the *Ae. tauschii* introgression population phenotyped under irrigated and non-irrigated conditions.

**Table S2.** Chromosome 6DL and 7DS haplotypes variants associated with spikelet number per spike and plant height in Ashland rainfed trial and their effects on other traits in the introgression population. Haplotype block HB1 - chr6D:463775852-463809722; haplotype block HB2 - chr7D:14185651-14596748; haplotype block HB3 - chr7D:14722457-14817138.
